# A realistic benchmark for the identification of differentially abundant taxa in (confounded) human microbiome studies

**DOI:** 10.1101/2022.05.09.491139

**Authors:** Jakob Wirbel, Morgan Essex, Sofia Kirke Forslund, Georg Zeller

## Abstract

**Background:** In microbiome disease association studies, it is a fundamental task to test which microbes differ in their abundance between groups. Yet, consensus on suitable or optimal statistical methods for differential abundance (DA) testing is lacking, and it remains unexplored how these cope with confounding. Previous DA benchmarks relying on simulated datasets did not quantitatively evaluate the similarity to real data, which undermines their recommendations.

**Results:** Here we develop a simulation framework which implants calibrated signals into real taxonomic profiles, including signals mimicking confounders. Using several whole-metagenome and 16S rRNA gene amplicon datasets, we validate that our simulated data resembles real data from disease association studies to a much greater extent than in previous benchmarks. With extensively parametrized simulations we benchmark the performance of eighteen DA methods and further evaluate the best ones on confounded simulations. Only linear models, *limma*, fastANCOM, and the Wilcoxon test properly control false discoveries at relatively high sensitivity. When additionally considering confounders, these issues are exacerbated, but we find that *post hoc* adjustment can effectively mitigate them. In a large cardiometabolic disease dataset, we showcase that failure to account for covariates such as medication causes spurious association in real-world applications.

**Conclusions:** For microbiome association studies tight error control is critical. The unsatisfactory performance of many DA methods and the persistent danger of unchecked confounding suggest these contribute to a lack of reproducibility among such studies. We have open-sourced our simulation and benchmarking software to foster a much-needed consolidation of statistical methodology for microbiome research.

## Introduction

The human gut microbiome is increasingly understood to play critical roles in host physiology and immunity, and thus mined for health and disease biomarkers. Taxonomic composition of gut microbial communities is highly variable between individuals^1,2^, yet clinical microbiome association studies strive to overcome inter-individual differences to identify microbial features that differ significantly between groups of individuals. Numerous diseases have been linked to specific microbes (including but not limited to inflammatory bowel diseases^3,4^, gastrointestinal cancers^5,6^, and cardiometabolic diseases^7^), typically by independently testing bacterial taxa for significant differential abundance (DA) between disease and control groups. DA methods loosely fall into three broad categories: a) classical statistical methods, b) methods adapted from (bulk) RNA-Seq analysis, or c) methods developed specifically for microbiome data.

While many studies have reported significant microbiome disease associations, some meta-analyses and cross-disease comparisons have suggested many of them to be unspecific or confounded^8–10^, i.e. attributable to other factors. For example, it is currently estimated that oral medication, stool quality, geography, and alcohol consumption collectively account for nearly 20% of the variance in taxonomic composition of gut microbiota^11,12^, yet these lifestyle factors often differ systematically between healthy and diseased populations being compared^13,14^. As a well-known example, two different studies reported associations between type 2 diabetes (T2D) and certain gut taxa which were later identified as a metformin response in a subset of T2D patients^8^. Furthermore, technical or batch effects resulting from non-standardized experimental protocols are prevalent in metagenomic studies^15,16^ and can outweigh biological differences of interest^6,17,18^.

Although unique characteristics of microbiome sequencing data – either whole meta-genome sequencing (WGS) or 16S ribosomal RNA amplicon sequencing (16S) – are well-described by now^19–21^, there is no consensus about the most appropriate DA procedures in the literature^22–29^. In principle, this is the purpose of benchmarking studies, which typically use parametric methods to simulate differentially abundant features under a ground truth for performance evaluation. Yet, for benchmarking conclusions to translate to real world applications, it is essential that simulated data recreate key characteristics of experimental data, which is why resampling techniques have also been used for benchmarking^26,30^.

Thus, on the more fundamental question of how best to simulate microbiome data, there is no consensus either. Existing simulation models have not been thoroughly validated on their ability to reproduce and resemble real experimental data (“biological realism”), nor has the impact of this property on downstream benchmarking applications been properly evaluated. Furthermore, despite a growing awareness of confounders in microbiome association studies, no evaluation of DA methods has meaningfully addressed this topic before.

To overcome these limitations, here we propose a simulation technique using *in silico* spike-ins into real data, causing specific taxa to differ in abundance and/or prevalence between two groups (imitating a case-control design), which we further extended to include confounding covariates with effect sizes resembling those in real studies. We quantitatively assess the degree to which parametric simulations employed in previous DA benchmarks lack biological realism, and show that the choice of simulation framework can explain divergent recommendations regarding DA methods. Based on our more realistic simulations, we perform a comprehensive benchmarking study of widely used DA methods and observe that many of these either do not properly control false positives or exhibit low sensitivity to detect true positive spike-ins. These issues were exacerbated under confounded conditions, but could largely be mitigated using the subset of methods which allowed post hoc adjustment for a covariate. Finally, we explore the merits of confounder-adjusted DA testing on a large clinical dataset.

## Results

### Assessment of biological realism for parametrically-simulated taxonomic profiles

As a first step, we aimed to explicitly evaluate how data generated from previous simulation frameworks compared to real metagenomic data. To do so, we simulated taxonomic profiles using the source code employed in previous benchmarks^23,25,26,31,32^, whereby case-control datasets were repeatedly generated with differentially abundant features of varying effect sizes (see **Methods**), and carried out a series of comparisons with the real input data. Simulation parameters were estimated in each case from the same baseline dataset of healthy adults, analyzed by shotgun metagenomic sequencing (Zeevi WGS)^33^. We observed the data simulated with every one of the previously used simulation models to be very different from real data as was apparent from principal coordinate analysis (see **Fig. 1a**) and from large discrepancies between the distribution of feature variances and sparsity (see **Fig. 1b**, **SFig. 1** and **SFig. 2**, also for other baseline datasets). Similarly, we observed the mean-variance relationships of many simulated features to fall outside the range of the real reference data, especially for the multinomial and negative binomial simulations (see **SFig 3a**). Finally we trained machine learning classifiers to distinguish between real and simulated samples (see **Methods** for implementation details), which was possible with almost perfect accuracy in nearly all cases, except for data generated from sparseDOSSA^31^ (**Fig. 1c, SFig. 2**). This classification attempt was motivated by the fact that machine learning, commonly employed in association studies to detect biomarkers, is highly sensitive to subtle differences between groups that may remain undetected by ordination-based analyses^10^. Overall, all of the assessed simulation frameworks produced unrealistic metagenomic data.

**Figure 1:**
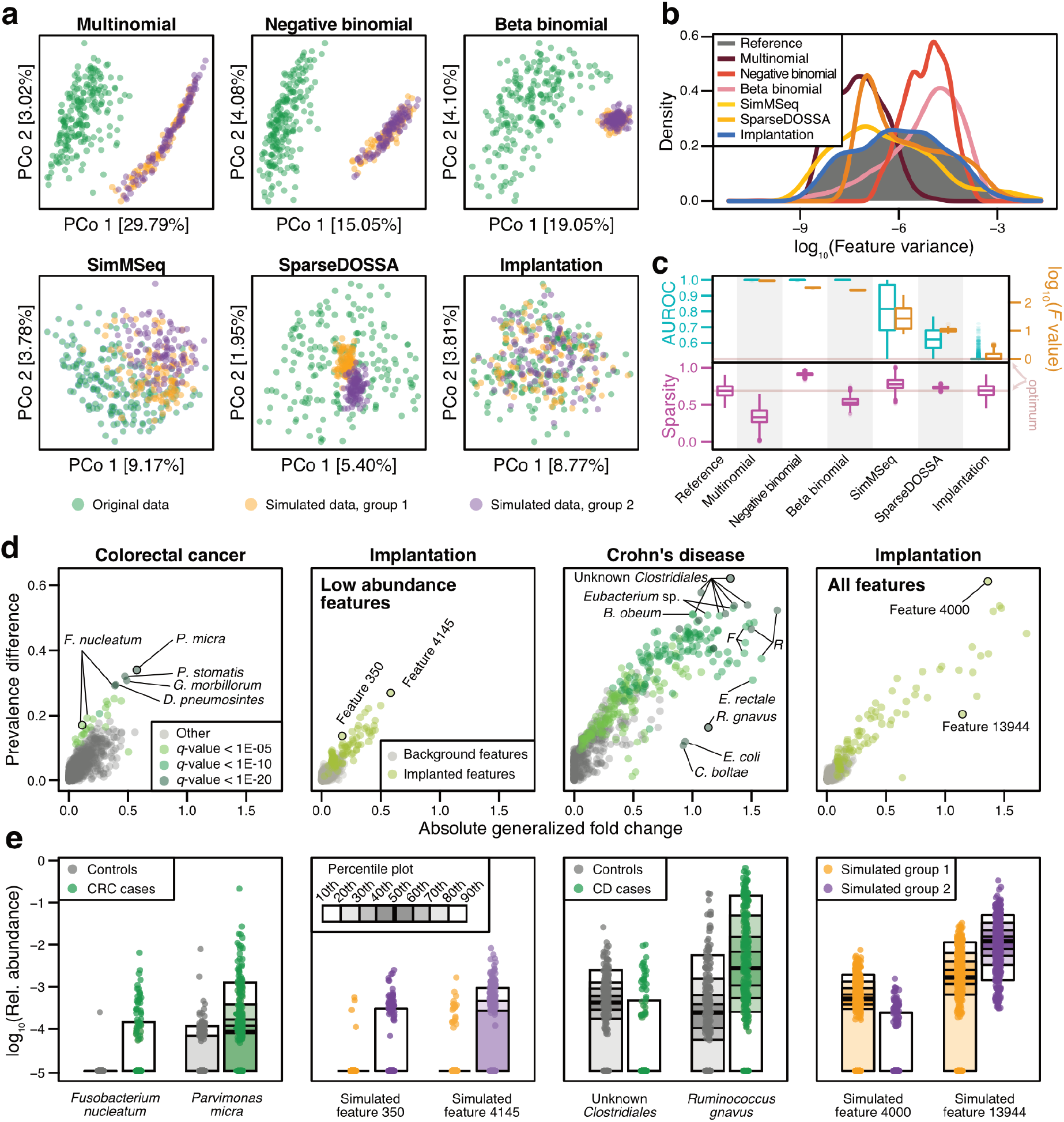
Signal implantation, but not parametric simulations, can reproduce key characteristics of metagenomic data and realistic disease effects. **a)** Principal coordinate projections on log-Euclidean distances for real samples (from Zeevi *et al*.^33^, which served as a baseline data set) and representative samples of data simulated in a case-control setting (group 1 and 2) using different simulation frameworks or signal implantation. For each method, the results from a single repetition and a fixed effect size are shown (abundance scaling factor of 2 with additional prevalence shift of 0.2 in our simulations, see **Methods** and **SFig. 4** for the complete parameter space). **b)** Distributions of log-transformed feature variances shown for the real and simulated data from a). **c)** The area under the receiver operating curve (AUROC) values from machine learning models (see **Methods**) to distinguish between real and simulated samples are shown across all simulated data sets in cyan. As complementary information, the log-transformed F values from PERMANOVA tests are shown in brown. Sparsity (fraction of taxa with zero abundance in a sample) is shown below in magenta. Boxes denote the interquartile range (IQR) across all values with the median as a thick black line and the whiskers extending up to the most extreme points within 1.5-fold IQR. **d)** The absolute generalized fold change^6^ and the absolute difference in prevalence across groups is shown for all features in colorectal cancer (CRC) and Crohn’s disease (CD). As a comparison, the same values are displayed for two data sets simulated using signal implantation (abundance scaling factor of 2, prevalence shift of 0.2), with implantations either into any feature or only low-abundance features (see **Methods**). Well-described disease-associated features are highlighted (*F*: *Faecalibacterium*, *R: Ruminococcus*) and selected bacterial taxa and simulated features are shown as percentile plot in e).

### Signal implantation yields realistic taxonomic profiles for benchmarking

To devise a more realistic simulation framework, we opted to manipulate real baseline data as little as possible by implanting a known signal with pre-defined effect size into a small number of features using randomly selected groups (similar to the downsampling approach of Jonsson et al.^30^, see **Methods**). As a baseline dataset, we chose the same study population consisting of healthy adults (Zeevi WGS)^33^, into which we repeatedly implanted a signal of differential abundance by multiplying the counts in one group with a constant (abundance scaling) and/or by shuffling a certain percentage of non-zero entries across groups (prevalence shift, see **Methods**). The main advantage of this proposed signal implantation approach is that it generates a clearly defined ground truth of DA features (like parametric methods) while retaining key characteristics of real data (a strength of resampling-based methods). In particular, feature variance distributions and sparsity were preserved (see **Fig. 1b, SFig. 1** and **SFig. 2**), as were the mean-variance relationships present in the real reference data (see **SFig 3a**), except at the most extreme effect sizes (see **SFig. 3b**). Consequently, neither principal coordinate analysis (**Fig. 1a**), nor machine learning classification could distinguish our simulated from the real reference data (**Fig. 1c, SFig. 2**). We observed these trends across all baseline datasets that we used for implantation (see **SFigs. 1-3**, see also **Methods** for details about included datasets).

### Implanted DA features are similar to real-world disease effects

To ensure our implanted DA features were comparable to those observed in real microbiome data in terms of their effect sizes, we focused on two diseases with well-established microbiome alterations, namely colorectal cancer (CRC)^5,6^ and Crohn’s disease (CD)^3,4^. In two separate meta-analyses (see **Methods**), we calculated generalized fold changes as well as the differences in prevalence between controls and the respective cases for each microbial feature (see **Fig. 1de**). The effect sizes in CRC were generally found to be much lower than in CD, consistent with machine learning results in both diseases (mean AUROC for case-control classification: 0.92 in CD and 0.81 in CRC, see ref^10^). For instance, the well-described CRC marker *Fusobacterium nucleatum* exhibits only moderately increased abundance in CRC, but strongly increased prevalence. This observation, generalizable to many other established microbial disease biomarkers, motivated the inclusion of the prevalence shift as an additional type of effect size for the proposed implantation framework. Depending on the type and strength of effect size used to implant DA features, the simulated datasets included effects that closely resembled those observed in the CRC and CD case-control datasets (**Fig. 1de**). In particular, simulated abundance shifts with a scaling factor of less than 10 were the most realistic and therefore used for subsequent analyses (**SFig. 4**).

### Performance evaluation of differential abundance testing methods

To benchmark the performance of widely-used DA testing methods under verified realistic conditions, eighteen published DA tests (see **Methods** for a list) were applied across all simulated datasets. Different sample sizes were created by repeatedly selecting random subsets from each simulated group, and each test was applied to the exact same sets of samples (see **Methods**). For each method, we used its recommended normalization and also explored additional data preprocessing techniques such as rarefaction (see **SFig. 5**).

The resulting *P* values were adjusted for multiple hypothesis testing with the Benjamini-Hochberg (BH) procedure to estimate method-specific false discovery rate (FDR) and recall; for each method, we then calculated the empirical FDR against the ground truth of our benchmarking data as the fraction of background features among all discoveries (using a cutoff of 5% estimated FDR). Additionally, a receiver operating characteristic (ROC) analysis was carried out to evaluate how accurately the raw *P* values could distinguish between ground truth and background features. In the ideal case of *P* values for all ground truth features being smaller than for any of the background features, the area under the ROC curve (AUROC) will be one; for random *P* values an AUROC of 0.5 is expected.

We found that for several methods the empirical FDR calculated against our ground truth far exceeded the estimated FDR (adjusted to 5%), especially for sample sizes under 200 (displayed for a single representative effect size in **Fig. 2a-c**, see **SFig. 6** for other effect sizes). In the most extreme case, and in line with previous reports^23,25^, the *fitZig* method from metagenomeSeq (*mgs*) displayed an empirical FDR of 80% (only 20% of features identified as significantly differentially abundant between groups were true *in silico* spike-ins). This behavior was observed across many sample and effect sizes (see **SFig. 6**). To explore whether this issue was due to the BH FDR estimation procedure, we additionally applied the more conservative Benjamini-Yekutieli (BY) method and found the empirical FDR to decrease, albeit at a loss of sensitivity (see **SFig. 7**). Nevertheless, for some methods, large discrepancies between empirical and estimated FDR were observed under both procedures (see **SFig. 7**), indicative of method-inherent lack of type I error control and in line with previous reports^26^. Of those DA methods for which the estimated FDR (resulting from BH correction) did not strongly deviate from the empirical FDR (i.e. empirical FDR not exceeding 20% in more than a fifth of settings between sample sizes 50 to 200), many also exhibited comparably low empirical recall or AUROC values (for example, *ANCOM* with a mean AUROC of 0.53 across all repetitions, see **Fig. 2d**), indicating these methods were relatively insensitive, too. In contrast, the methods with the highest AUROC values were the linear models (*limma*, and the general *LM*), the *Wilcoxon* test, and *fastANCOM*, which also exhibited proper FDR control at comparably high sensitivity (**Fig. 2**). These methods ranked at the top with remarkable consistency across other datasets (**Fig. 2d**), thereby emerging as reliable testing frameworks for the analysis of human-associated microbiome data.

**Figure 2:**
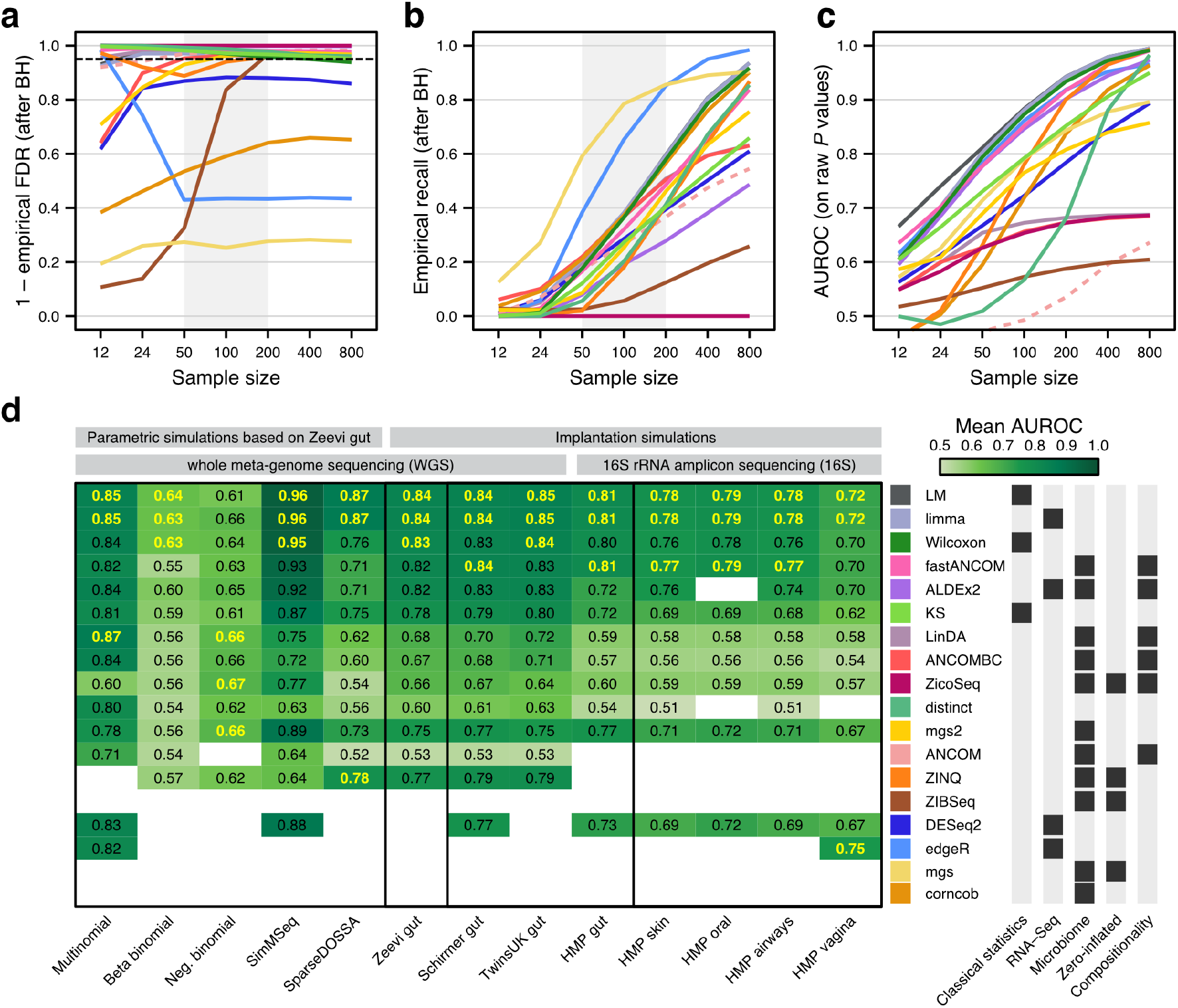
Performance evaluation of differential abundance testing methods and simulation strategies. For a signal implantation simulation with a single, moderate effect size combination (abundance scaling factor of 2, prevalence shift of 0.2, all features eligible for implantation), **a)** the mean empirical FDR and **b)** the mean empirical recall (calculated after Benjamini-Hochberg (BH) correction of raw *P* values) are shown for all included DA test methods across different sample sizes (see **Methods**). Additionally, mean AUROC values for differentiating between implanted and background features (calculated from raw *P* values) are shown in **c)**. The nominally expected value of a 5% FDR is indicated by a dotted black line. Since ANCOM does not return *P* values (see **Methods**), empirical FDR and recall were based on the recommended cutoffs (without adjustment) and therefore highlighted by dashed lines. **d)** The mean AUROC values across all effect sizes and repetitions for the sample sizes 50, 100, and 200 (shaded area in **a**-**c**) are depicted in the heatmap for the different simulation strategies and baseline datasets, including non-gut human-associated microbiomes. The methods are ordered by their AUROC values on the Zeevi WGS gut dataset, with methods that exceed a mean empirical FDR of 10% in more than 20% of settings ranked after those that do not. The three top-performing methods ranked per column are highlighted in yellow. AUROC values are not shown if the mean empirical FDR exceeded 20% (four times the nominal value) in more than 20% of settings.

### Impact of simulation approach and method normalization choices on benchmarking results

Our benchmarking results did, however, depend on the simulation method used for data generation. Negative binomial simulations in particular ranked LinDA, ZicoSeq, and metagenomeSeq as the top DA methods, almost entirely discordant with the rankings from other simulations, datasets, and biomes (see **SFig. 2d** and **SFig. 8**). For both the negative and beta binomial simulations, no DA method (including the respective top three) had a mean AUROC > 0.7 across sample sizes 50-200 (i.e. selection criteria for **Fig. 2d**). Interestingly, of the DA methods which had the overall poorest performance (i.e. exceeded 20% false discoveries across at least 20% of simulations), four of the bottom five assume one of these distributions in their models.

Most of the included DA methods use count tables as input and might perform method-specific normalization or variance-stabilizing procedures (see **Methods**). To explore the effect of rarefaction, one of the most commonly employed library size normalization techniques in microbiome data analysis, we additionally ran all methods with rarefied counts as input. On average, rarefaction led to a reduction of sensitivity (lower empirical recall and lower AUROC values) across the majority of tested methods without improving the empirical FDR (see **SFig. 5**). For methods that do not model count data, we also explored other commonly used data preprocessing techniques, such as the total sum scaling (TSS) or centered log ratio (clr) transformation (see **Methods** for details). With these, recall and AUROC of *limma* and the *LM* considerably improved after TSS-log preprocessing, whereas the performance of the *Wilcoxon* test decreased after application of the clr and robust clr transformations (see **SFig. 5** and **Methods**).

While it is not disputed that sequencing produces compositional (i.e. proportional) data^19^, it is unclear from the literature to which extent this is problematic in downstream analysis and whether computational approaches can successfully address it^34^. Advocates of such approaches argue that (unaddressed) compositionality leads to spurious associations as a consequence of the inherent correlation structure between features. To minimize unintended compositional correlation structures in our simulation framework, signal implantation alternated between groups by default (following Weiss *et al.*^25^, see **Methods**). However, to assess DA method behavior in the presence of strong compositionality, we conducted an additional simulation with intentional compositional effects by implanting signals into only one group (see **Methods**), which led to abundance shifts in background features (i.e. spurious signals) in addition to the ground truth signals (see **SFig. 9**). Under these conditions, the empirical FDR increased for all tested methods with increasing effect sizes, with *ANCOM* and *ALDEx2* being least affected (see **SFig. 9**).

### DA method performance evaluated under confounded conditions

One major issue with the simple DA testing procedure outlined above is that it does not consider potentially confounding covariates (i.e. variables tracking the myriad ways case-control groups may differ in addition to disease status) as an important source of spurious associations. To mitigate this issue, association studies increasingly rely on *post hoc* stratification or multiple regression techniques to adjust for potential confounders in DA testing, but how well this works for microbiome studies has not yet been quantitatively evaluated.

To close this gap, we considered two common scenarios encountered in real studies, reflecting (i) a disease signal of interest appearing stronger in one cohort of a multi-center study (i.e. technical confounding), or (ii) the disease group in a case-control study taking medication while the control group does not (i.e. biological confounding). Technical confounders tend to manifest broadly across many features^35^, while most commonly administered medication impact fewer microbial taxa or act more narrowly^36^. Given a case-control study (with case-control status represented by a binary variable) where a confounding covariate can be represented by another binary variable, a dataset can then be divided into four groups. When scenario (i) or (ii) is applicable, i.e. when these two variables are associated with one another (as captured by the phi coefficient, see **Methods**), groups are imbalanced with respect to one another in the sense that most samples are observed in only two of the four theoretical groups. In order to invoke both scenarios, we implemented a resampling algorithm that generates testing data according to a prespecified phi (i.e. confounder strength, see **Fig. 3a** and **Methods**), and then we examined the prevalence of both types of confounders in real datasets to ensure the validity of our chosen parameters (see **SFig. 10a** and **SFig. 11b** for (i) and (ii), respectively).

**Figure 3:**
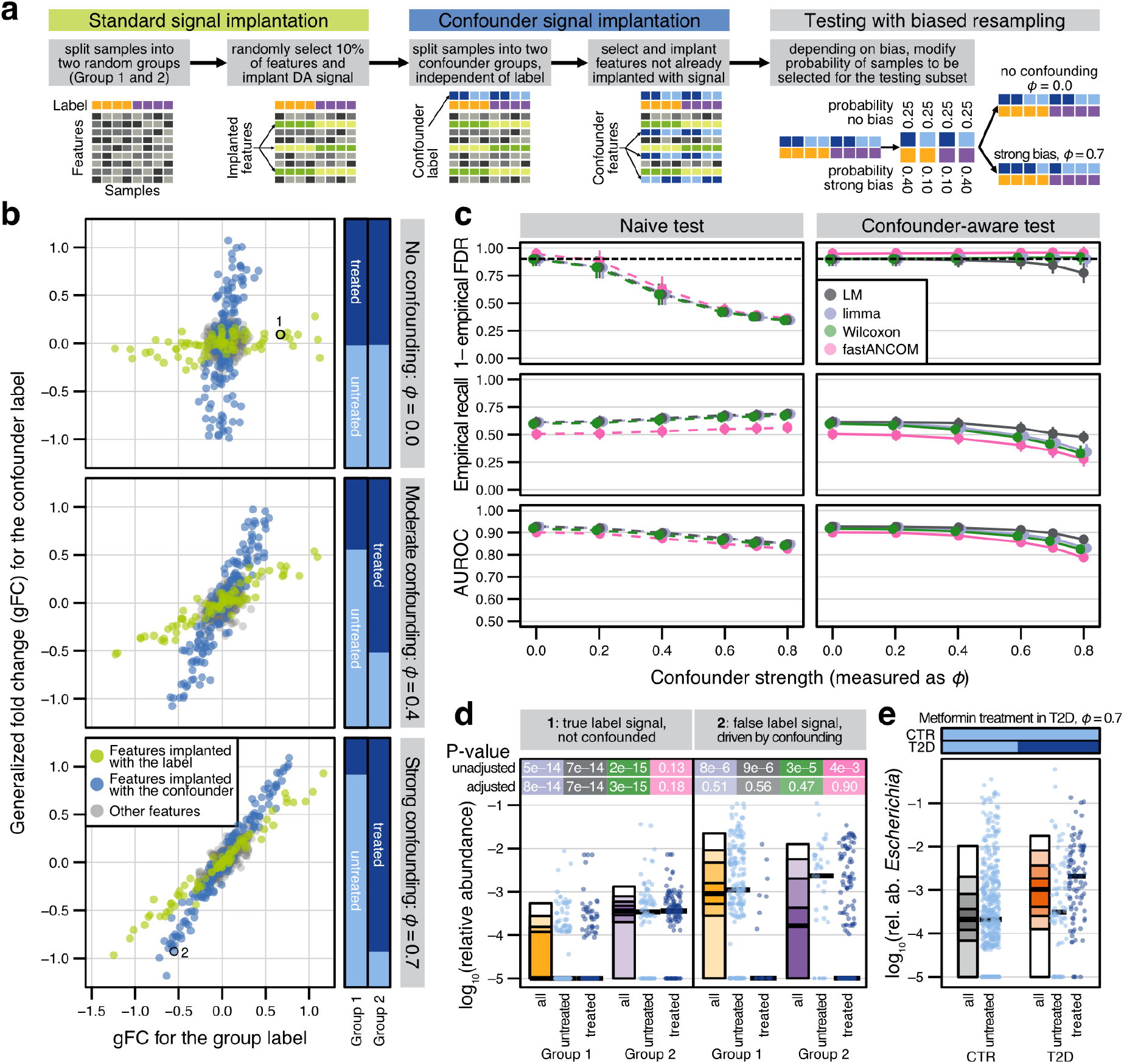
Loss of precision and recall under confounding can be alleviated by *post hoc* adjustment. **a)** Using a single dataset, DA features were independently implanted into a small proportion of taxa for both a main group label (as described above) and for an independent binary (confounder) label, imitating e.g. disease and medication status labels, respectively. Subsets for DA testing were generated using a parameterized resampling technique such that the degree of association between these two variables could be modified. **b)** Generalized fold change (gFC) calculated for the label is contrasted to the gFC calculated for differences between confounder values across all bacterial taxa (abundance scaling factor of 2, prevalence shift of 0.2, all features eligible for implantation, a single representative repeat shown). Bars at the right visualize the confounder strength by showing the proportion of confounder-positive samples in each group (with ϕ=0 serving as unconfounded control). Main implanted features are highlighted in green and features implanted for the confounder label are in blue. **c)** Mean empirical FDR, empirical recall (both calculated after BH-correction) and AUROC (on raw *P* values) for sample size 200 and the same effect sizes as shown in a) were computed for tested DA methods, using unadjusted and confounder-adjusted test configurations. Error bars indicate standard deviation around the mean for all repeats. **d)** Simulated (log_10_ relative) abundances plotted by main and confounder labels (See Fig. 1 for definition of abundance quantiles), with both unadjusted and confounder-adjusted significance shown at top, colored as in c). **e)** *Escherichia* abundance appears naively associated with type 2 diabetes, yet is driven by metformin intake in a subset of diabetics (reproduced from Forslund *et al.*^8^)

To first create benchmarking data that resembled confounding due to study heterogeneity, we combined samples from pairs of real population studies in healthy adults^33,37,38^ and used study origin as the covariate in our biased resampling approach (see **Methods**). For the evaluation of DA methods, we focused on those found to control the FDR at concomitant high sensitivity in our previous benchmark and which could additionally be adjusted for covariates (*limma*, the *LM*, *fastANCOM,* and the *Wilcoxon* test). We compared these methods in their naive (unadjusted) and adjusted configurations on our simulated data.

In the negative control setting, both studies were equally sampled, such that the study variable was not associated with the main grouping variable (see **SFig. 12**). Here, expectedly, naive and adjusted performances were similar to the previous (unconfounded) benchmark results. However, greater confounder strengths, simulated by disproportional sampling of diseased cases from one study and controls from another, resulted in increased false discoveries and overall poor performance of the naive models, which was partially but not fully rectified in the adjusted models (see **SFig. 12**).

Next, we considered confounding due to medication akin to cases described in the literature^8,13,14^. To generate benchmarking data with narrower confounding effects, we modified our implantation procedure to include an additional binary grouping variable associated with a distinct set of spiked-in DA features, and again applied our biased resampling approach (see **Methods** and **Fig. 3a**). The negative control setting (generated by even sampling producing four balanced groups, which can occur for e.g. matched demographic factors like sex), manifested as non-overlapping effects between the two variables (see **Fig. 3b**). In contrast, larger bias values (representing e.g. a higher prevalence of medication intake in the disease group) resulted in progressively overlapping groups, thereby making it difficult to attribute differential abundance in individual taxa to the respective grouping variables (i.e. to distinguish disease from medication effects, see **Fig. 3b**).

We found that in this scenario naive testing led to dramatically inflated empirical FDR as confounder strength increased (empirical FDR ∼40% at moderate confounding with ϕ=0.4, see **Fig. 3c**). Adjusted tests, however, generally maintained FDR control, although performance of the *LM* weakened under extreme confounding (implemented as a linear

mixed-effect model, *LMEM*, per the discussion in Gomes^39^, see **Methods** and **SFig. 13**). The *LM* nonetheless exhibited the highest overall AUROC, results which were consistently observed in other baseline datasets and at varying proportions of implanted confounder-associated features (see **SFig. 14**).

To illustrate method behavior at the level of individual simulated feature abundances, we selected true and spurious signals from the ground truth, under a control setting and under strong confounding, respectively (see **Fig. 3b**). In the control setting, only *fastANCOM* failed to identify the true DA feature (see **Fig. 3d**, panel 1), which was not entirely unexpected considering its lower overall sensitivity (see **Fig. 3c**). In contrast, when biased resampling produced an association between the two grouping variables, we observed a clear false positive driven by confounding, as could be diagnosed from greatly decreased statistical significance when comparing the adjusted with the naive DA test results (see **Fig. 3d**, panel 2).

Importantly, our bias parameterization (confounder strength) in these simulations tracked well with phi coefficients calculated on real data (see **SFig. 11**), and our implanted effect sizes resembled the actual effects observed for metformin treatment in T2D patients (see **Fig. 3e**). Overall, our results suggest that measured confounders (as long as ϕ<0.8, see **Fig. 3e**) can be effectively controlled or adjusted for during *post hoc* statistical testing.

### Discerning robust from confounded associations in real datasets

To further explore the consequences of naive association testing compared to *post hoc* adjustment, we applied both approaches to gut metagenomic samples from cardiometabolic disease patients in the MetaCardis cohort^14,40,41^. The strongest confounding potential was seen for chronic coronary artery disease (CCAD) and commonly-indicated medications taken by a large fraction of these patients, especially statins and aspirin (ϕ=0.89 and ϕ=0.9, respectively), as well as type 2 diabetes (T2D) and metformin (ϕ=0.72, see **SFig. 11**). Four linear models were built for each disease-drug combination (naively testing for disease or drug associations, respectively, and corresponding adjusted models for each) across all species-level taxonomic abundances. The resulting coefficients and *P* values were used to classify each feature with respect to both drug and disease associations (see **Methods**). As expected, large phi coefficients manifested as a strong linear relationship between naive *LM* coefficients, confirming that confounding can be diagnosed from such models (compare **Fig. 3b** and **Fig. 4a**). Importantly, the inclusion of random effects in the adjusted models helped to disentangle these overlapping effects and exposed drug- or disease-specificity in numerous individual associations (**Fig. 4a**).

**Figure 4:**
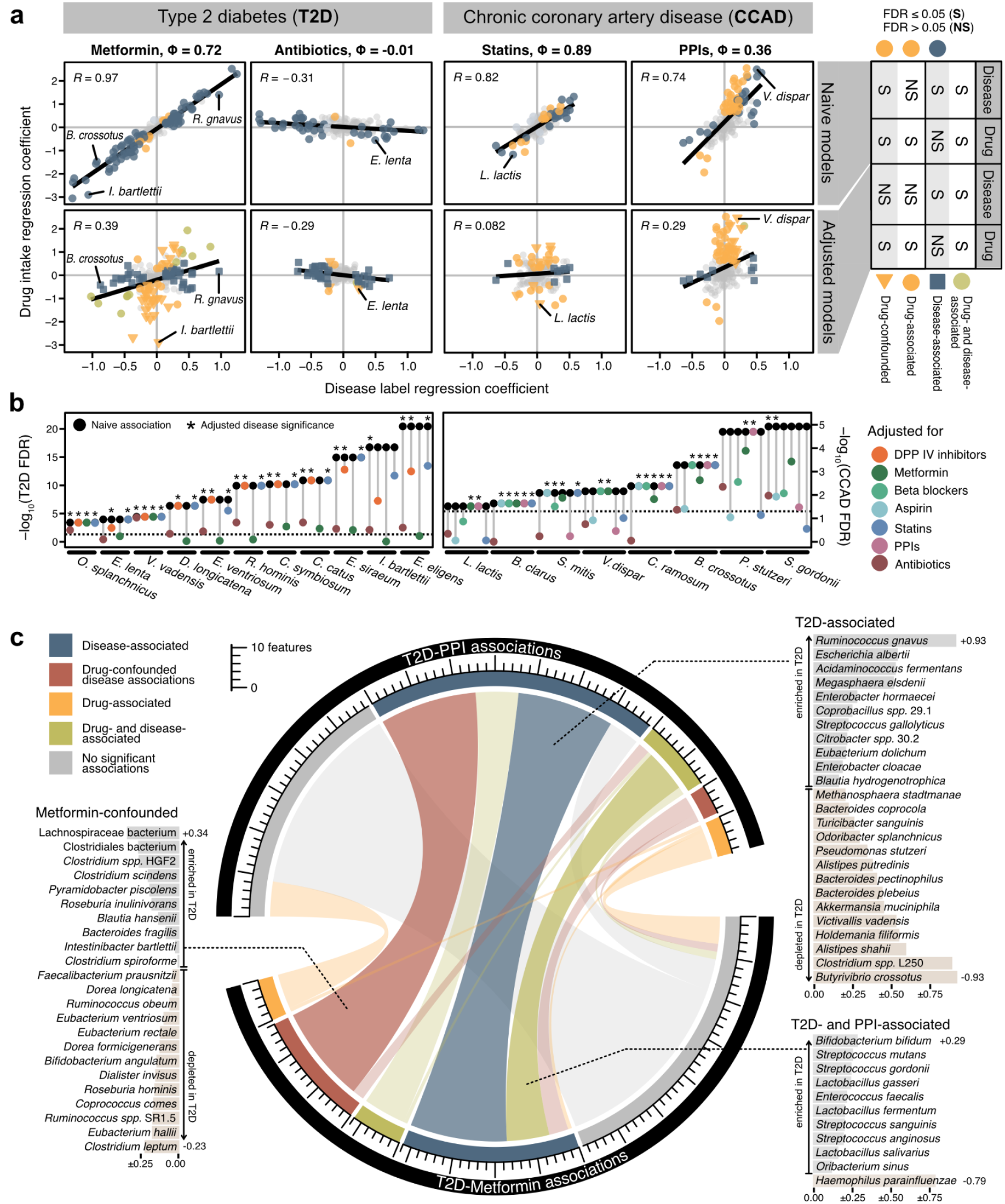
Linear models are capable of disentangling drug- and disease-associated microbial features. **a)** Regression coefficients from a subset of disease-drug combinations comparing naive linear models to adjusted mixed-effect models for all bacterial taxa. Adjusted models included a second term (either drug intake or disease status for x and y axes, respectively) as a random effect, which diminished the strong linear dependence between naive model coefficients (shown). When the significance of each term was compared between the naive and adjusted models (see **Methods**) drug-specific or confounded effects were revealed in some features. **b)** Exemplary subset of features displaying either the largest number of significant disease associations across different drug-adjusted models or the largest reductions in disease coefficient significance upon adjustment (i.e. most confounded). **c)** Comparison of feature classifications (see **Methods**) from the metformin- and PPI-adjusted disease association models across all bacterial taxa. Integrating information across models restricts disease associations to a more robust subset and reveals drug-confounded associations. Adjusted T2D regression coefficients are shown in light gray or light brown bars behind species names (indicating enrichment in T2D or control group, respectively).

In contrast to CCAD (and other diseases), T2D exhibited larger, more significant naive associations with more taxa (**Fig. 4b**). A little less than half of these were confounded by metformin treatment, as identified by the loss of a significant association with T2D and retention of a significant association with metformin in the adjusted models (**Fig. 4b** and **Methods**). Adjusting for antibiotic intake, on the other hand, did not significantly reduce the number of T2D-associated taxa, but generally reduced their coefficient size and significance. CCAD-associated taxa were sensitive to adjustment by multiple different drugs, including antibiotics, probably reflecting the complexity and variation typically seen in disease association studies where medication records are thoroughly analyzed. In general, confounder-adjusted linear models effectively helped to disentangle disease- and drug-associated taxa (evident from a reduced number of significant disease associations), consistent with their demonstrated ability to distinguish true positives from confounder positives in our simulation benchmarks (see **Fig. 3c**).

Metformin and proton pump inhibitors (PPIs) were among the largest drug effects observed in our analysis. Whereas most metformin-associated taxa were also naively T2D-associated, most PPI-associated taxa were not. This is in line with previous reports demonstrating metformin intake to correlate with T2D disease severity^8,14,42^, and PPI-associated gut microbiota changes as a disease-independent presence of mainly oral commensals^43,44^ (consistent with the drugs’ mechanism of action). To further tease out disease-associated taxa, we cross-referenced the feature classifications we obtained from our models across different drugs as a robustness analysis. For example, PPI-adjusted models resulted in disease associations, several of which were metformin-confounded. By integrating information across both sets of confounder-adjusted models, we could uncover a more robust subset of disease-associated taxa (see **Fig. 4c**). Taken together, this analysis suggests that linear mixed-effect models are an effective and versatile method to *post hoc* improve robustness of findings in association studies in which medication records are available.

## Discussion

Clinical microbiome research typically involves differential abundance testing to detect associations between host phenotypes, including many human diseases or responses to treatment^45^, and individual microbial features in high-throughput. Numerous DA methods have been developed, encompassing a broad range of assumptions and hypotheses, but their performance has remained controversial, several previous benchmarking studies notwithstanding^22–27^. Based on our results, we argue that the failure to produce consensus is partially explained by their reliance on unvalidated and ultimately unrealistic parametric simulations of microbiome data as ground truth. Here, we addressed this by implanting minimal, biologically-motivated modifications into real taxonomic profiles. Additionally, we extended our simulation framework to incorporate effects resembling confounders frequently encountered in microbiome studies. Although our strategy may lack the theoretical appeals of a parametric mathematical model, we empirically verified that our simulated taxonomic profiles were the only ones to retain essential metagenomic data properties and produce data that are virtually indistinguishable from real samples. Furthermore, our framework still provides the flexibility to specify effect and sample sizes needed for an extensive evaluation against a ground truth, which was not the case in previous benchmarks built upon real datasets^26,27,46^.

To make empirically-guided recommendations on the suitability of DA methods, we performed a neutral benchmark based on our implantation framework that included the most widely used DA methods. Evaluating each DA test on nearly one million simulated case-control data sets, we found that most methods yielded an excess of false positives especially on smaller sample sizes (N<100, see **Fig. 1**). Notable exceptions were the non-parametric *Wilcoxon* test, *limma*, *fastANCOM,* and *LMs*, for all of which we found the empirical FDR to be close to theoretical expectations while high sensitivity was retained across a range of sample and effect sizes, datasets, and human-associated microbiomes.

Unlike previous benchmarks^23,30^, our results suggest DA methods borrowed from RNA-seq analysis, with the exception of *limma*, perform poorly on taxonomic microbiome profiles (further discussed in **Supplementary Notes 1** and **2)**. Surprisingly, we also found most methods developed specifically for microbiome data to have comparably low power and high false positives rates (with the exception of *metagenomeSeq2*) across the range of dataset sizes most commonly seen in contemporary studies (see **Fig. 2d**). On a positive note, at least when when applied to larger samples (N ≥ 200 per group), more DA tests (including both *ANCOM* and *ANCOM-BC*, *ZIBseq*, *ZINQ*, and *metagenomeSeq2*) controlled the empirical FDR at the nominal level. Overall, our findings strongly support the use of classical statistical methods (linear models or the nonparametric *Wilcoxon* test) or the recently developed *fastANCOM,* and suggest that many studies employing other methods may have reported a substantial fraction of spurious microbiome-disease associations.

This inferential risk is further exacerbated by confounding factors, which are rarely adjusted for in microbiome disease-association studies, if recorded at all. While medication can be an obvious confounder for disease associations^14^, one study based on the American Gut Project dataset identified various lifestyle and physiological parameters, for example alcohol intake or stool quality, as additional sources of heterogeneity^13^. As a more straightforward and scalable alternative to the solution proposed in that study (matching for all potential confounders in resampled groups), we explored simpler *post hoc* adjustment in *LMs*, *limma*, *fastANCOM,* and the *Wilcoxon* test using simulated data which mimicked well-understood^8,12–14,36^ confounders. Reassuringly, adjusting the respective DA tests restored unconfounded performance to a large extent.

Limitations of our work include a narrow focus on human-associated taxonomic profiles from cross-sectional study designs, which precludes generalization to non-human microbial communities and longitudinal study designs. Moreover, we have not examined biologically-relevant associations with e.g. strain-level functional traits^47^, or host-microbe rhythmicity^48^, and we only minimally explored differential sequencing depth confounding in our simulations, which has been investigated in more detail elsewhere^32^. Additionally, our confounded simulations were restricted to exploring the impact of a single binary variable. Yet, linear (mixed-effect) models are very flexible and computationally efficient tools, which can readily accommodate continuous covariates such as stool microbial load^34,49^ and account for repeated measurements^29^. Notably, most newer compositional methods we tested (i.e. ZicoSeq^32^, *ANCOM-BC*^50^, *LinDA*^51^, and *fastANCOM*^52^) adjust for a derived bias correction factor in a similar linear regression framework, in contrast to earlier methods relying on pairwise log-ratios (ANCOM^53^) or permutations on transformed values (ALDEx2^54^), both of which were prohibitively computationally intensive (see **SFig. 15**).

In our benchmark, we implemented simple *LMEM*s from the *lmerTest* R package^55^. Statistically analogous implementations to ours are available through *MaAsLin2*^56^, which also includes several count-based linear models, and *SIAMCAT*^10^, which provides dedicated functionality to check for potential confounders in the metadata before employing several confounder-adjusted DA association tests. In our analysis of drug effects in T2D, we demonstrated the importance of further integrating deconfounded information to reveal robust disease-associated feature subsets. Analogous logic may be found in the vibration-of-effects paradigm^57,58^ and the *metadeconfoundR*^14,59^ package, which screens for potential confounders before performing combinatorial nested model DA tests on naive and confounder-adjusted linear models (from the *lme4* package^60^) to classify feature robustness. Given the limiting prerequisite that covariates need to be recorded for explicit adjustment, methods that account for broadly-manifesting unmeasured variables (as with population structure in genome-wide association studies^61,62^) could help to minimize confounding variation in microbiome studies as well. As more attention is paid to the complexity of factors at play in clinical microbiome studies, the need for DA tools to not only accommodate potential confounders in their association tests, but also to automate high-throughput robustness checks, is increasingly apparent.

In our view, the unsatisfactory performance of a wide range of DA methods and the persistent danger of unchecked confounding in the literature warrant a community effort to develop and benchmark more robust methodology. To assist researchers in developing and validating new DA methods, or establishing further benchmarks, both our signal implantation framework and our benchmarking analysis were designed to be easily extensible and are available as open source code (see **Methods**). Ultimately, community-driven benchmarking efforts similar to DREAM challenges^63^ or the Critical Assessment of Metagenome Interpretation (CAMI)^64^ project could accelerate the much-needed consolidation of statistical methodology for microbiome research.

## Methods

The first part of this Methods section describes the design of our simulations and its comparison to previous approaches as well as real microbiome sequencing data. The code for generating the simulated data is made available in an R package called SIMBA (https://github.com/zellerlab/SIMBA).

The second part of this Methods section contains details on how various differential abundance testing methods (in combination with different preprocessing routines) were applied to the simulated data, and how their results were evaluated against the ground truth and visually compared using custom scripts provided through the BAMBI R project (https://github.com/zellerlab/BAMBI).

### Data selection and preprocessing

The dataset from Zeevi *et al*.^33^ was used as a baseline for the simulations in the main text. We included only those samples that had been analyzed via whole metagenome sequencing (WGS) and removed samples analyzed by 16S ribosomal RNA amplicon sequencing (16S). Additionally, we used the datasets from Schirmer e*t al*.^38^ (WGS) and Xie *et al.*^37^ (TwinsUK WGS) as independent baselines for the simulation framework evaluation and assessment of microbiome data properties (see **SFig. 1-3**), and combined different combinations of two WGS datasets at a time to mimic study effects in a meta-analysis setting (see **SFig. 12**). All three WGS datasets consist of human gut microbiome samples. To explore other human-associated microbiomes, we also ran SIMBA on the abundance tables from the HMP1 dataset^65,66^, which included samples from different body sites. HMP 16S abundance tables and metadata were downloaded from the data portal at https://portal.hmpdacc.org/.

For the gut WGS datasets, raw data were downloaded from ENA (PRJEB11532 for Zeevi, ERP010708 for TwinsUK, and PRJNA319574 for Schirmer) and analyzed as described before^10^. In short, after preprocessing and removal of host contamination, taxonomic profiling was performed with mOTUs2 (v2.5^67^). All input data tables were filtered within SIMBA for prevalence (at least 5% across the complete dataset) and abundance (maximum relative abundance across samples of at least 1e-04). In the case of repeated samples per patient, we selected only the first time point for each patient.

Lastly, the MetaCardis dataset^14^ was used to explore drug confounding in real data. Cell-count adjusted, quantitative mOTUs profiles and metadata from the publication were downloaded from Zenodo (https://doi.org/10.5281/zenodo.6242715); a pseudocount of 1 was applied to zero counts before log transformation. In the case of repeated samples per patient, we selected only the first time point for each patient.

### Parametric methods for the simulation of metagenomic data

To simulate metagenomic data on the basis of parametric methods, the implementations described in previous differential abundance benchmarking efforts were adapted into SIMBA (re-using the authors’ original source code wherever possible) and are briefly summarized here:

● Both McMurdie and Holmes^23^ and Weiss *et al*.^25^ used multinomial-generated counts, but differed slightly in how they created differentially abundant features. If not indicated otherwise, results for multinomial simulations were based on the implementation from Weiss *et al*., since the resulting effect sizes were closer to real effects (see **SFig. 4**).
● Hawinkel *et al.*^26^ included two different univariate parametric simulations based on the negative binomial and the beta binomial, as well as the multivariate Dirichlet distributions, all of which were included in SIMBA. Their non-parametric and real data shuffling methods were excluded on the grounds that they lacked sufficient parameterization for downstream benchmarking. For the beta binomial and optionally the negative binomial distribution, the correlation structure between bacterial taxa was estimated using SPIEC-EASI^20^ as in the original publication. Values generated from the Dirichlet distribution were converted into counts via a post-processing rounding step.
● The Bayesian semiparametric method from Yang and Chen^32^ was reproduced in SIMBA via the SimMSeq function of the GUniFrac R package.
● Lastly, to simulate data as described in Ma *et al.*^31^, SIMBA relied on the dedicated functions in the sparseDOSSA R package.

Differentially abundant features were introduced into each parametric simulation as described in the respective original publications. For the multinomial simulations from McMurdie and Holmes as well as for the sparseDOSSA approach, features were scaled in abundance after the simulation was completed. In the case of the other simulation methods, the underlying parameters were adjusted with a scaling factor before the simulation. A range of effect sizes (abundances scaled by multipliers of 1, 1.25, 1.5, 2, 5, 10, and 20) was explored and for each effect size, a total of 20 repetitions was simulated (for each simulation method). At an abundance scaling factor of 1, no effects were introduced into the data and therefore those repeats served as internal negative controls. Simulation method implementations can be found in the respective helper_xxx.R files in SIMBA, and scripts to automate data generation on a SLURM cluster are stored in the create_simulations folder of the BAMBI repository.

### Implantation framework for the realistic simulation of microbiome data

To create benchmarking datasets without a parametric model, we implemented a novel simulation framework referred to as signal implantation (helper_resampling.R in SIMBA). In each simulated repetition, the original samples were randomly split into two groups and 10% of features were randomly selected to become differentially abundant between the groups. Effects were implanted both via scaling abundances (using the same effect sizes as the parametric simulations) and by shifting prevalences (0.0, 0.1, 0.2, and 0.3).

For the abundance scaling, count values in one group were multiplied with a scaling factor to increase the abundances. The prevalence shifts were implemented by identifying non-zero counts in one group and randomly exchanging a specific percentage of those with occurrences of zero abundances in the other group (if possible), thereby creating a difference in prevalence across the groups with respect to the selected feature. The implantation of signals alternated between the two groups in order to prevent a systematic difference in total count number across groups (inspired by the considerations in Weiss *et al*.^25^). For each combination of effect sizes (abundance scaling and prevalence shift), 100 repetitions were simulated for the dataset from Zeevi *et al.* and 20 repetitions for all other datasets.

Knowing that some diseases preferentially associate with high- or low-abundance taxa (as in e.g. colorectal cancer, see **Fig. 1de**), we additionally explored criteria to determine the set of features eligible for signal implantation: namely, *all* - all taxa were equally likely to be selected to carry a signal, or *low* - only low abundance features (the 75th percentile across all samples not exceeding 0). Other criteria we explored yielded unrealistic effect sizes (see **SFig. 4**), and were not pursued further.

As a last step, the resulting generalized fold change^6^ between the groups for all implanted features was recorded. Features with a fold change lower than 0.001 (resulting mostly from low-prevalence features being selected for implantation) were rejected and not recorded as implanted signals (this resulted in the removal of zero to three features across all repetitions of each simulated effect size, on average, depending on effect size).

To generate simulations that mimic compositional effects, the signal implantation was carried out as described with the modification that signals were implanted into one group only (not alternating between groups). Then, the number of counts for each sample was scaled down to the original value of the unaltered sample by rarefaction using the *vegan* R package^68^.

### Reality assessment for simulated data

To evaluate how well simulated metagenomic data approximated real data, for each “group” in a simulation file, we calculated the sample sparsity, feature variance and mean together with differences in prevalence and the generalized fold change^6^ between mock groups. Additionally, the separation between original and simulated samples in principal coordinate space was evaluated using PERMANOVA as implemented in the *vegan* package^68^. As a complementary approach, a LASSO logistic regression machine learning model was trained to discriminate between real and simulated samples using the SIAMCAT R package^10^ and the AUROC of the cross-validated model was recorded. In short, real and simulated data were combined into a single feature table, with the label for classification being either “real” or “simulated” (independent of the groups used for signal implantation). After log-standardization of relative abundances, a LASSO model was trained with a ten-times repeated ten-fold cross-validation scheme and the average performance recorded.

### Included DA testing methods

To evaluate the performance of various DA testing methods, the R implementation of each method was incorporated into SIMBA using the recommended normalizations, if applicable, as described below (with default parameters if not stated otherwise). The following methods were included in the benchmark (methods which allowed for confounder-adjustment by inclusion of covariates into the model and were assessed here are denoted with an asterisk*), usually available through an R package of the same name (version and installation routes are in the *renv.lock file* of the BAMBI repository):

● *\*Wilcoxon*: for the naive Wilcoxon test, the *wilcox.test* function available through base R was used per taxon; per default in R, the ranks of tied observations were averaged. For confounder-adjusted testing, the *wilcox_test* function of the *coin* package^69^ was used with formula “*feature ∼ label | confounder”*.
● Kolmogorov-Smirnov test (*KS*): the *ks.test* function available through base R was used per taxon.
● *Linear models (*LM*): for naive testing with linear models, the function *lm* available through the base R distribution was used per taxon. *P* values were then extracted by applying the function *anova* on the trained model. For the confounder-adjusted testing, the confounder variable was included as a random effect in the model formula using the *lmer* function of the *lmerTest* package^55^ (formula “*feature ∼ label + (1|confounder)*”). The main grouping variable (label) was tested for significance using the base R *summary* function on the fitted *lmerModel* object. We also tested different linear model formulae, with little difference in performance (see **SFig. 13**).
● *\*Limma*^70^: the *lmFit* function was used with the complete feature matrix and the label as design matrix as input. *P* values were then extracted after applying the *eBayes* function on the resulting *MArrayLM* object, which applies a moderated t-statistic to the linear model coefficients. For confounder-aware testing, the confounder was supplied to the *lmFit* function call as *block* parameter.
● *edgeR*^71^: normalization factors were estimated using the *calcNormFactors* function with the *RLE* method across the whole dataset, as in Nearing *et al*^72^. Then, the *estimateCommonDisp* and *estimateTagwiseDisp* functions were applied to the *DGEList* object before differential abundance testing was performed with the *exactTest* function (also in the package).
● *DESeq2*^73^: as recommended in the *phyloseq* vignette^74^, the geometric mean for each sample was added to the *DESeqDataSet* object as a normalization factor. Finally, differential abundance was calculated with the function *DESeq* which uses the *nbinomWald* test function (also in the package) as the default.
● *ALDEx2*^54^: the *aldex* function was used with default parameters, which specify a Welch’s t-test on log-transformed and centered data.
● *mgs* and *mgs2* (*metagenomeSeq*^22^): low prevalence features (<5% across all samples) and samples with fewer than ten counts were filtered out. As recommended in the *metagenomeSeq* vignette, a normalization factor was calculated for each sample via the *cumNormStat* function and added to the *MRExperiment* object. For testing, two different models can be fitted within the same R package, which are included here as *mgs* (using the *fitZig* function) and *mgs2* (using the *fitFeatureModel* function), analogously to Weiss *et al*.^25^
● *ZIBSeq*^75^: the *ZIBSeq* function in the package with the same name was used. The included option to perform a *sqrt* method-specific normalization was run separately (*ZIBSeq_sqrt*).
● *Corncob*^76^: the feature matrix was transformed into a *phyloseq* object and the *differentialTest* function was applied to test each feature (using the formula *“∼ label”*). Per the method default and suggestion in the vignette, the Wald test was used for hypothesis testing.
● *ZINQ*^77^: the *ZINQ_tests* function was applied to each bacterial taxon, and the resulting *P* values were calculated with the *ZINQ_combination* function using default parameters.
● *distinct*^78^: the *distinct_test* function was used with default parameters.
● *ANCOM*^53^: no dedicated R package is available from the original publication and the standard implementation is prohibitively slow for larger benchmarks (see **SFig. 15**). Therefore, we used the implementation available through Lin *et al*.^79^. Since *ANCOM* does not return *P* values, its primary outputs (W values) were converted into a score ranging between 0 and 1 for easier evaluation through the same framework as the other methods. The recommended decision threshold for discoveries in *ANCOM* is equal to 0.7 x number of tested taxa. Therefore, the W values above this decision threshold were transformed into scores lower than 0.05 (corresponding to a ‘discovery’ in our evaluations), whereas all other W values were monotonically transformed to range between 0.05 and 1. Since this score does not constitute real *P* values, but rather a convenience for scoring the output of the package with the same functionality as the output of other methods, we did not apply any multiple hypothesis correction on the transformed *ANCOM* score.
● *ANCOM-BC*^50^: the *ancombc function from the ANCOMBC package was used. The lib_cut* parameter (indicating a minimum number of counts per sample) was set to 100.In contrast to ANCOM, ANCOM-BC outputs *P* values.
● *fastANCOM*^52^: the *fastANCOM* function from the package with the same name was used with default parameters. For confounder-adjusted testing, the confounder variable was passed as parameter *Z* to the *fastANCOM* function.
● *LinDA*^51^: the *linda* function in the *LinDA* package was used with the following parameters, in concordance with the GitHub README: *lib_cut = 1000, prev.cut = 0.1, windsor.quan = 0.97* and raw *P*-values were extracted out of the resulting list.
● *ZicoSeq*^32^: the *ZicoSeq* function with default parameters was used (based on the *GUniFrac* package vignette), except for the following: top-end Winsorization (*winsor.end = ’top’*), low prevalence filter (*prev.filter = 0.1, max.abund.filter = 0.002*), and square-root transformation (*link.func = list(function (x) x^^^0.5)*, as shown in the vignette of the package). The raw *P*-values were extracted and FDR-corrected for all subsequent analyses. *ZicoSeq* also provides a specialized permutation-based FDR control, which was not assessed in this work due to its computational burden.

### Additional preprocessing transformations

Most included DA methods take read count values as input and either work on those directly or perform specific normalizations, which are described above (and evaluated in **SFig. 5**). To explore the effect of a preprocessing step commonly applied in microbiome data analysis, we (optionally) included *rarefaction* of counts as an additional transformation (before method-specific normalizations were applied) for all methods. This transformation consisted of downsampling counts to the 25th percentile of the total counts across samples.

The DA methods *Wilcoxon* test, *KS* test, *LM*, and *limma*, do not explicitly model count data and work with a variety of data distributions. Therefore, we applied a range of other transformations that have been widely used in the microbiome field. Overall, the applied transformations consisted of: *clr* (centered log ratio transform), *rclr* (robust centered log ratio transform), *TSS* (total sum scaling), *TSS.log* (total sum scaling, followed by log_10_ transformation of the data), and *TSS*.*arcsin* (total sum scaling, followed by the arcsine square root transformation), *rarefaction-TSS* (rarefaction followed by total sum scaling), and *rarefaction-TSS.log* (rarefaction followed by total sum scaling and log_10_ transformation of the data).

### Benchmarking of DA testing methods at different sample sizes

To simulate different sample sizes, we randomly selected *n* samples out of the two groups (*n*/2 from each) for each combination of effect size and each repetition. These samples were saved via indices such that for comparisons each method was applied to the exact same data. Seven different sample sizes were explored (12, 24, 50, 100, 200, 400, and 800) and 50 sets of test indices were created for each. For the evaluation of a single DA method, a total of 980,000 unique configurations were generated and used as input (7 abundance shifts x 4 prevalence shifts x 100 implantation repeats x 7 sample sizes x 50 subsamples as testing repeats).

The *P* values across all taxa were recorded and adjusted for multiple hypothesis testing using the Benjamini-Hochberg procedure^80^. If no *P* value was returned for a specific taxon (because the taxon had been filtered out by a method-specific filtering step, for example), we set this value to 1 instead before BH adjustment. To evaluate the performance of each method, we checked the raw *P* values from each testing scenario for how well bacterial taxa with differential abundance were detected, calculating an AUROC score using the raw *P* values as a predictor. The empirical FDR and recall were calculated using the BH-corrected *P* values at a cutoff of 0.05.

### Generating confounders through biased resampling

We identified two high-risk confounding scenarios relevant to real clinical microbiome studies^5,6,8,14^, representing both biological and technical factors known to influence community composition, and extended our framework in order to simulate data for benchmarking under both scenarios.

First, samples were given an additional binary confounder label and features were chosen for implantation on the basis thereof. For the “narrow” confounding simulations (mimicking e.g. medication intake in a case-control disease study and impacting few features only), an additional set of unique features was implanted to differ between the groups implied by the confounder label as described above, resulting in two distinct sets of ground truth features. Alternatively, for the “broad” confounding simulations (mimicking e.g. batch effects in a meta-analysis and impacting a majority of features) in which we combined two independent datasets at a time, this confounder label denoted the bonafide study membership (Zeevi^33^, TwinsUK^37^, or Schirmer^38^ depending on the combination, see **SFig. 12**). In both scenarios, four distinct groups of samples were created.

Second, resampling was parameterized in order to calibrate the proportions of samples drawn from each of the four groups in the final testing subsets, thereby modulating the association between the main group and confounder variables and generating a confounded signal, as defined and measured by the extent of overlap between the implanted signals (see **SFig. 11**). We encode the confounder strength as the “bias” term and it is analogous to the phi coefficient as a measure of association between two binary variables. While phi is a standardized metric ranging from -1 to 1 (with 0 indicating no association and ±1 indicating perfect positive or negative association, respectively), our confounder strengths functionally range from 0 to 1, corresponding to observations in real data where a control group is present (see **SFig. 11**). At a bias value of 0, all four groups will be proportionally represented in each resampled testing subset and hence there is no confounding. Larger bias values will correspond to non-uniform representation of confounder labels among testing subsets, which will create a correlation structure between the signals implanted according to the confounder label and the groups to be differentiated by the DA methods, mimicking confounding in real case-control studies.

### Effect size assessment in real case-control datasets

To compare simulated data to real case-control microbiome studies, we collected datasets for two diseases with a well-described microbiome signal. For colorectal cancer (CRC), we included the data from five studies^6,81–84^ conducted across three continents, which were the basis for an earlier meta-analysis that identified consistent microbial biomarkers for CRC^6^. For Crohn’s disease (CD), we similarly included five case-control studies^4,85–88^ that had been analyzed previously^10^. For CD, the data were restricted to the first measurement for each individual, whenever applicable. The data from all studies were taxonomically profiled via mOTUs2^67^ (v2.5) and features were filtered for at least 5% prevalence in at least three of the studies. Differences in prevalence across groups and the generalized fold change were calculated for each microbial feature as previously described^6^ and the significance of enrichment was calculated using the blocked *Wilcoxon* test from the *coin* package in R^89^.

### Confounder and robustness analysis in the MetaCardis data

To deepen our understanding of confounder effects found in real clinical microbiome data, we evaluated a subset of drug-disease combinations from the MetaCardis cohort, which were preprocessed as described above. Each disease subcohort was combined with the control group to constitute a case-control dataset, and the phi coefficient was calculated using a custom implementation of the standard formula^90^ with respect to each binary drug intake metadata variable (see **SFig. 11a**). For each bacterial taxon, two naive linear models were built using the base R *lm* function which modeled bacterial abundance as a function of either disease status or drug intake only, and two corresponding confounder-adjusted models were built using the *lmer* function from the *lmerTest* package^55^, and additionally incorporated drug intake or disease status as a random effect, respectively. Significances of the resulting coefficients were adjusted for multiple testing according to the Benjamini-Hochberg procedure^80^ and used to classify associations (at an FDR of 0.05).

For a given disease-drug combination, taxa which were significantly associated with the disease status and the drug intake in all four models were assigned a “drug- and disease-associated” status. Taxa bearing a significant disease association in both naive and adjusted models which did *not* possess a significant association with drug intake in the adjusted models were classified as “disease-associated”. Lastly, taxa which were significantly associated with both drug and disease in the naive models, but no longer with disease in the adjusted models, were considered to be “drug-confounded”. Taxa which had no significant associations with disease but significant drug associations in both naive and adjusted models were simply “drug-associated.”

### Implementation

The codebase for the presented results is split into two projects. The first one, an R package called SIMBA (<u>S</u>imulation of <u>M</u>etagenomic data with <u>B</u>iological <u>A</u>ccuracy, available at https://github.com/zellerlab/SIMBA), provides the modular functionality to (i) simulate metagenomic data for a benchmarking project, (ii) perform reality checks on the simulated data, (iii) run differential abundance (DA) testing methods, and finally (iv) evaluate the results of the tests. The second project, BAMBI (<u>B</u>enchmarking <u>A</u>nalysis of <u>M</u>icro<u>B</u>iome Inference methods, available at https://github.com/zellerlab/BAMBI), is a collection of R scripts relying on the *batchtools* package^91^ in order to automate and parallelize the execution of SIMBA functions. Both SIMBA and BAMBI are available through GitHub and will enable other researchers to explore a similar benchmarking setting for other baseline datasets, other biomes, and additional DA testing methods. As part of the respective GitHub repositories, we included vignettes to showcase the functionality with toy examples. For reproducibility and to allow direct comparison of new methods with those in the presented benchmark, the simulation files and statistical results presented in this manuscript are available on Zenodo (see **Data Availability**). The *renv* package manager^92^ was used to install all software and to document versions, and our computing environment may be instantiated on new machines using the *renv.lock* file in the BAMBI repository.

### Data structures

To efficiently store and organize the large amount of related data required to evaluate both metagenomic simulation and differential abundance methods, we designed SIMBA around the Hierarchical Data Format (HDF5)^93^. Although we opted to work with R (v4.0.0) and the *rhdf5* package, HDF5 files are language-independent.

To ensure that the exact same input data was used for each DA test, we implemented our framework to pass a specific set of normalized feature vectors (e.g. bacterial taxon counts) to any implemented method. The samples to be included in each test are stored via their indices in the HDF5 format; for example, when testing on sample sizes n=100 and n=200 for 50 iterations, there would be a 50x100 and a 50x200 matrix of sample indices stored for each effect size parameterization of each simulated dataset.

## Author contributions

G.Z. and S.K.F. conceived the study and supervised the work. J.W. and M.E. implemented the software based on a prototype by G.Z. and performed the statistical analyses with guidance from G.Z. and S.K.F.. J.W., M.E., and G.Z. designed the figures with input from S.K.F.. J.W., M.E., and G.Z. wrote the manuscript with contributions from S.K.F. All authors discussed and approved the final manuscript.

## Code availability

The software package to simulate metagenomic data (SIMBA) is available on GitHub: https://github.com/zellerlab/SIMBA. Similarly, the repository containing the scripts to run a benchmark (BAMBI) is also available on GitHub: https://github.com/zellerlab/BAMBI.

## Data availability

The synthetic datasets used in this manuscript and source data for all figures are available at Zenodo (https://doi.org/10.5281/zenodo.8429303); this includes preprocessed species-level taxonomic profiles and metadata from all cohorts used in our analyses.

## Acknowledgements

We would like to thank members of the Zeller and Forslund groups for inspiring discussions, as well as Dr. Ami Bhatt and her group. We are indebted to the European Molecular Biology Laboratory (EMBL) for providing HPC resources as well as to its IT services team and Yan Ping Yuan for technical support. This work was partially funded by EMBL and MDC. G.Z was supported by the Federal Ministry of Education and Research (BMBF; the de. NBI network grant no. 031A537B and grant no. 031L0181A) and the German Research Foundation (DFG; CRC SFB-1371). S.K.F. was supported by the German Research Foundation (CRU KFO339, CRCs SFB-1365, SFB-1470 and SFB-1449) and the European Commission (HORIZON-HLTH-2022-STAYHLTH-02: IMMEDIATE).

