## Supplementary Material for "A realistic benchmark for the identification of differentially abundant taxa in (confounded) human microbiome studies"

### Supplementary Information

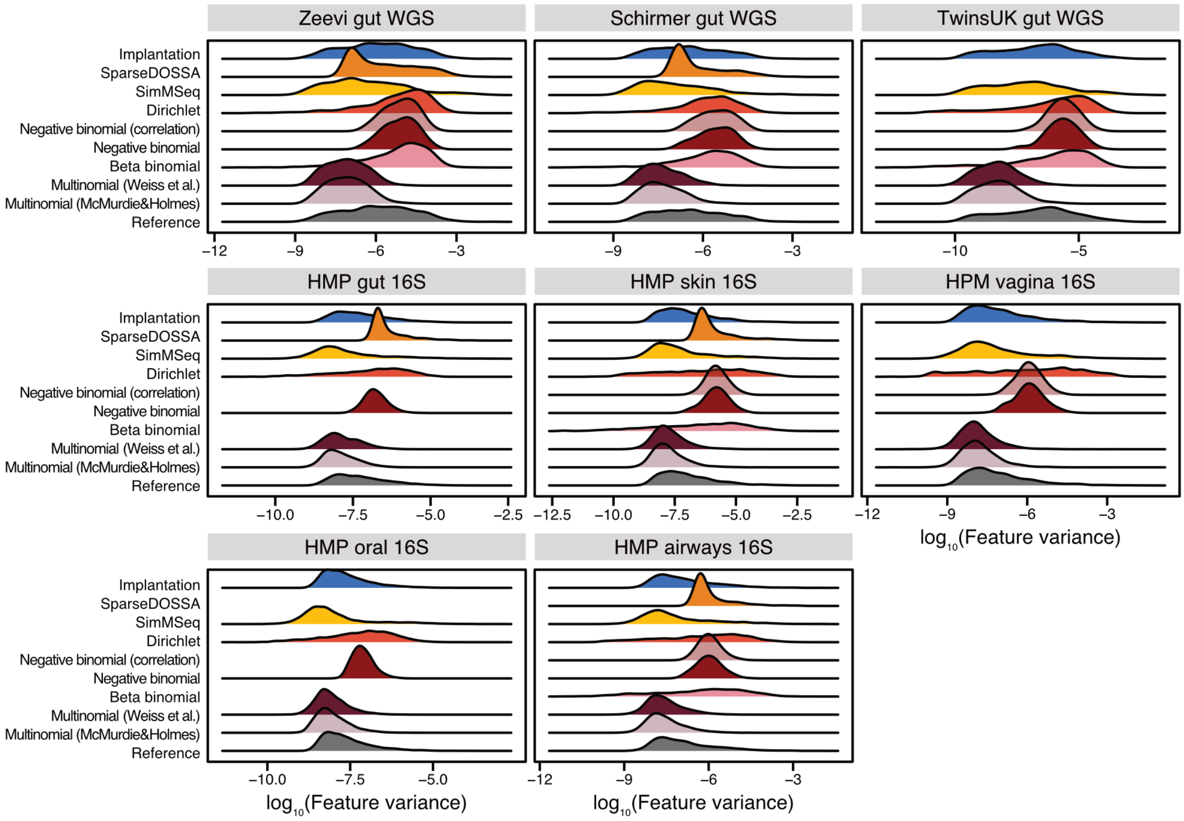


**Supplementary Figure 1: Distribution of feature variances is preserved in signal implantation, but not parametric simulations.** For all baseline datasets (see **Methods**), the distribution of feature variances were recorded for the real input data (reference, gray) as well as simulated data from different parametric simulation frameworks or the signal implantation framework. For simulated data, the distributions are shown for a single repeat of a fixed effect size (abundance scaling of 2, prevalence shift of 0.2, if applicable). Missing ridges for some datasets indicate the simulation procedures failing to converge after five days of compute time.

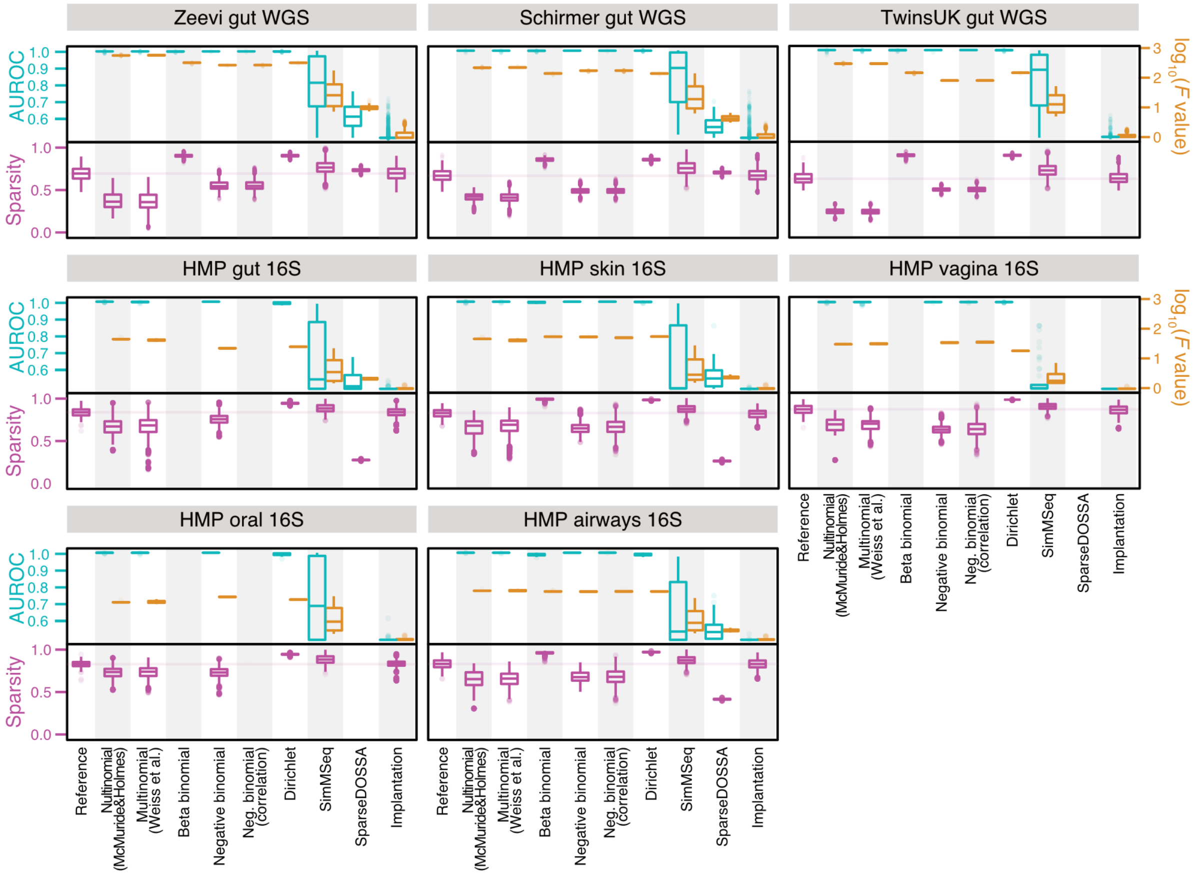


**Supplementary Figure 2: Sample sparsity, machine learning, and PERMANOVA analyses indicate that signal implantation, but not parametric simulations, can reproduce key data characteristics across various datasets.** For all baseline datasets (see **Methods**), sample sparsity was recorded as the fraction of zero-abundance features per sample for the real input data (reference) as well as data simulated with different parametric simulation frameworks or the implantation framework. The distribution of sample sparsity is shown as magenta box plots, with the median line of the reference extended across each subplot. Similarly, the AUROC values resulting from machine learning analyses (a measure for how well a machine learning model can distinguish between real and simulated samples, see **Methods**) and the log-transformed F-values from PERMANOVA analyses (a measure for how different real and simulated samples are from one another, based on log-Euclidean distances between samples) are shown as cyan and brown boxplots, respectively. Boxplots show the measures for all repetitions (100 for the dataset from Zeevi WGS and 20 for all other datasets) and effect sizes. Missing boxes for some datasets indicate the simulation procedures failing to converge after five days of compute time. See **Fig. 1** in the main text for box definitions.

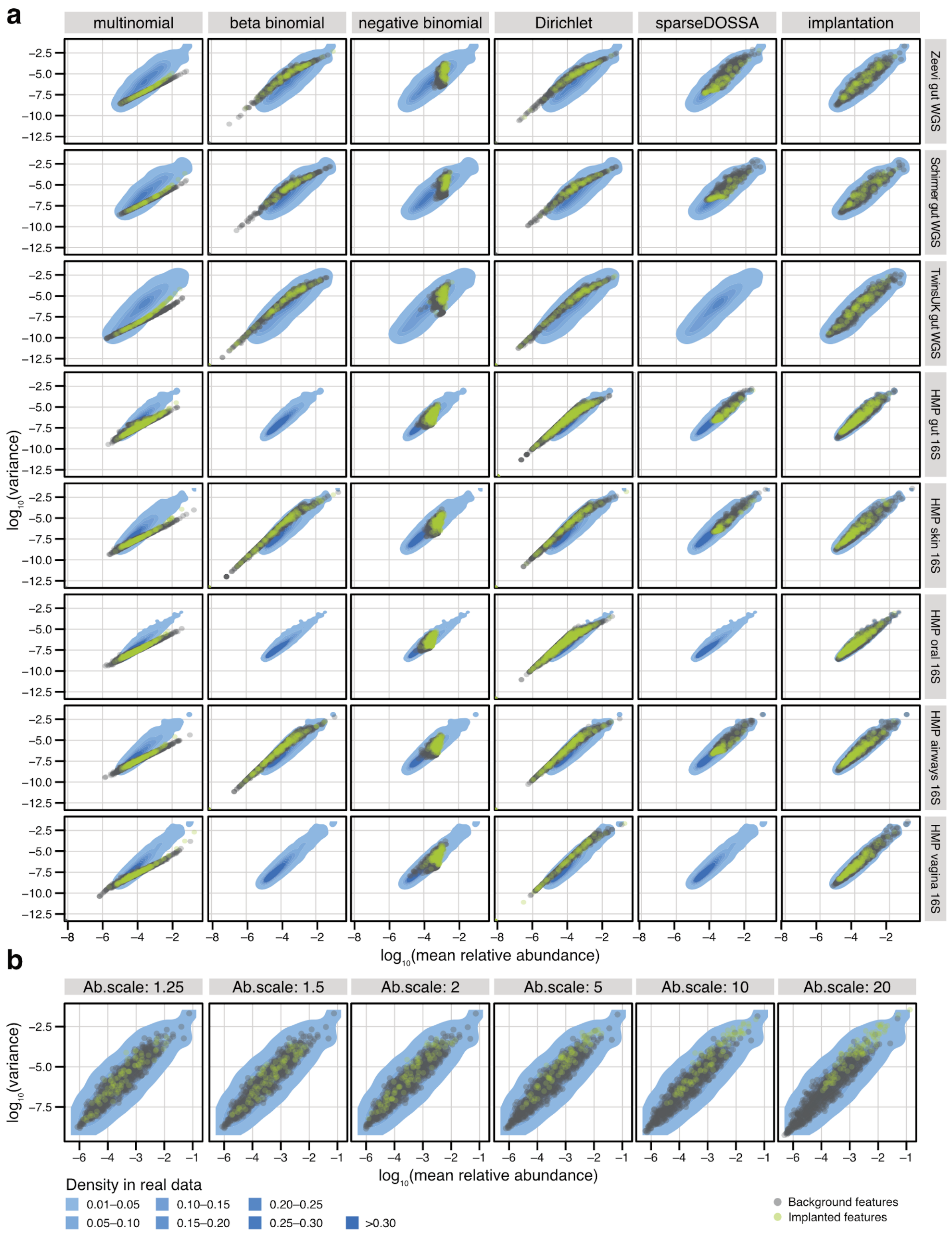


**Supplementary Figure 3: Mean-variance relationship is retained in data derived from signal implantation, but not parametric simulations. a)** For all baseline datasets, the log10-transformed mean relative abundance is plotted against the log10-transformed variance for all taxa simulated with five parametric simulation frameworks and the signal implantation framework. A single repetition of a fixed effect size (abundance scaling of 2, prevalence scaling of 0.2, if applicable) is shown. Features that have been selected for differential abundance are highlighted in green. The grey shaded area indicates the density of the mean-variance relationship in the real input data, estimated through the `MASS::kde2d()` function in R. Missing points for some datasets indicate that the simulation procedures failed to converge after five days of compute time. **b)** For the Zeevi gut WGS dataset, the mean-variance relationship is shown for various abundance scaling effect sizes (fixed prevalence scaling of

0.2, a single random repetition). Using the approach of the PERMDISP test (`vegan::betadisper`), the dispersion of real and simulated mean-variance values was tested for significant differences. For an abundance scaling factor of 10, 46% of repetitions resulted in significant ($P < 0.05$) differences in the dispersion between real and simulation data, depending on the implanted prevalence shift (prevalence shift of 0: 7%, pr. shift of 0.1: 40%, pr. shift of 0.2: 60%, and pr. shift of 0.3: 77% of repetitions), whereas 97% of repetitions for abundance scaling of 20 showed significant differences. None of the tests for other abundance scaling effect sizes were significant. All parametric simulation frameworks resulted in data with significantly different dispersion between real and simulated mean-variance values irrespective of the effect size used.

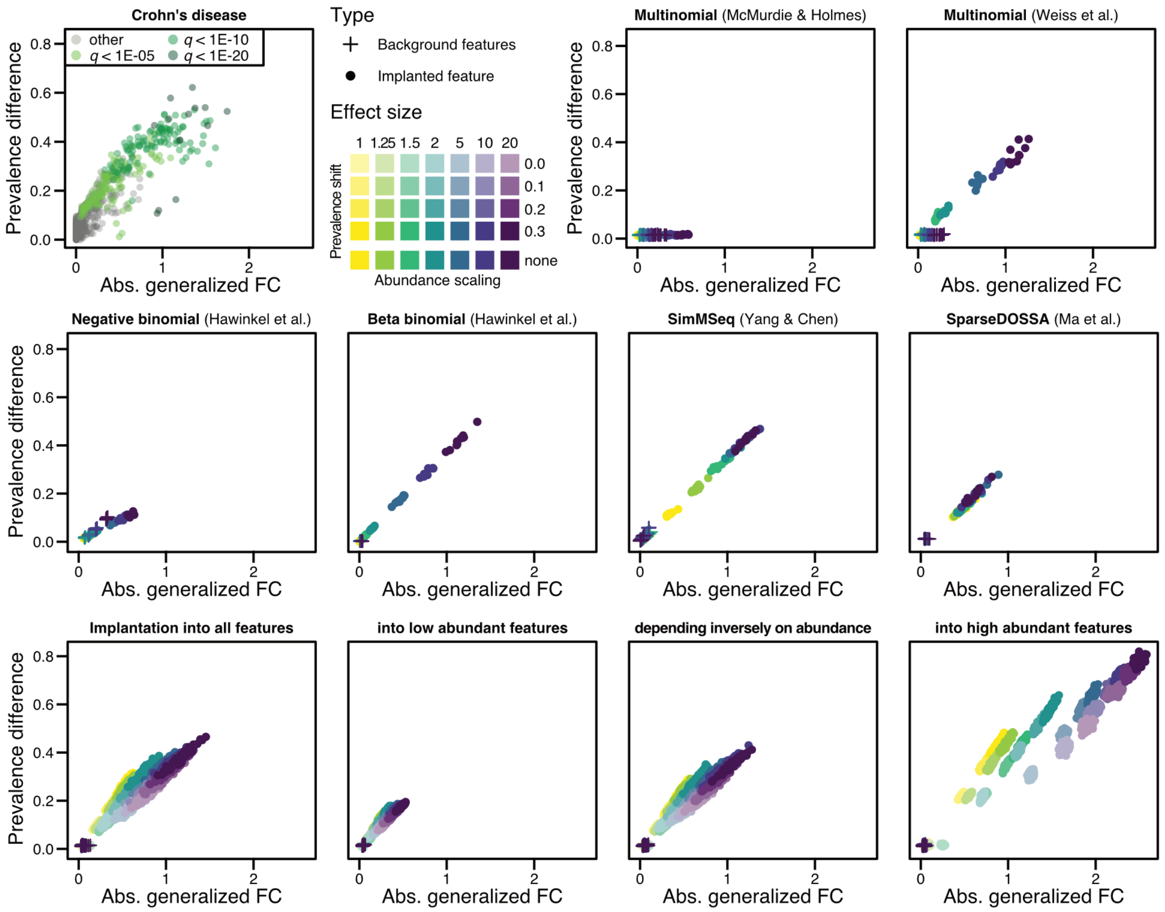


**Supplementary Figure 4: Implanted effect sizes vary across different simulation schemes and eligible feature sets.** The absolute generalized fold change (gFC)[1] and the absolute prevalence difference between groups was calculated for all features across all repetitions in every simulation scheme. For each repetition, the mean gFC and mean prevalence difference values were calculated for both background and implanted features (ground truth differentially abundant features). As a reference, the real gFC and prevalence shift values observed across all features in the Crohn's disease meta-analysis are shown in the top left panel (see **Methods** and main text **Fig. 1**, also for a comparison to gFCs and prevalence shifts observed in colorectal cancer). In the bottom row, mean gFC and mean prevalence shift values are shown for different signal implantation configurations that vary feature sets eligible for implantation (see **Methods**). Signals implanted into high abundant features resulted in (mean) effect sizes which were much larger and deviated substantially from real Crohn's disease data.

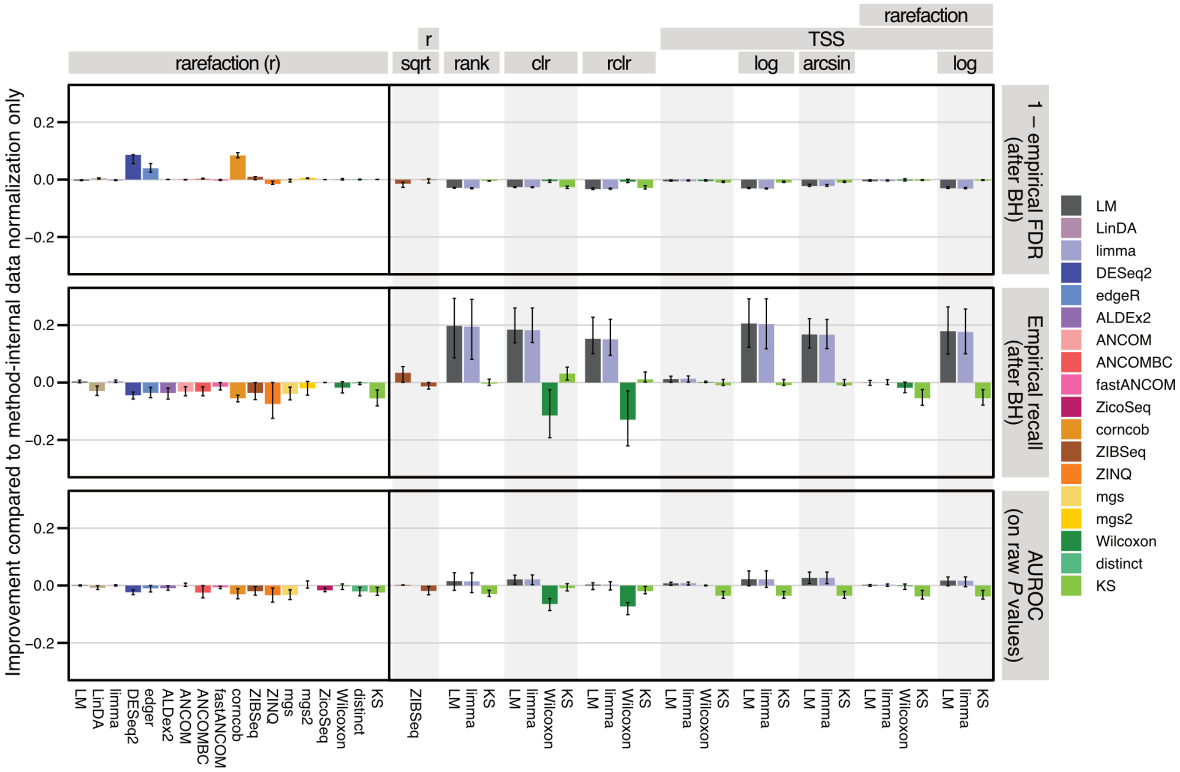


**Supplementary Figure 5: Performance of differential abundance testing methods tends to decrease after rarefaction.** The mean change in empirical FDR, empirical recall and AUROC for the detection of ground truth differentially abundant (DA) features are shown as bars across all tested effect and sample sizes to compare method-internal normalization (exclusively) with prior data transformation (assessing methods that are commonly used in microbiome data analysis, such as e.g. rarefaction). Negative values denote decreased performance when compared to no data preprocessing, and error bars indicate the standard deviation across all repetitions. The horizontal bars on the top of the plot indicate the preprocessing method used (and combinations thereof, e.g. total sum scaling (TSS) could be combined with rarefaction and/or other downstream transformations). For all DA testing methods, rarefaction was applied before any method-specific normalization was performed. Overall, rarefaction led to a decrease in empirical recall and lower AUROC scores across most methods. Other data preprocessing methods were applied for methods that do not specifically model count data and could therefore be supplied with other types of data. Both *limma* and the *LM* showed marked improvements in recall (with comparably low deterioration of empirical FDR) across several preprocessing techniques, where the compositional preprocessing methods *clr* and *rclr* lowered the AUROC and the recall for detection of ground truth DA features for the *Wilcoxon* test.

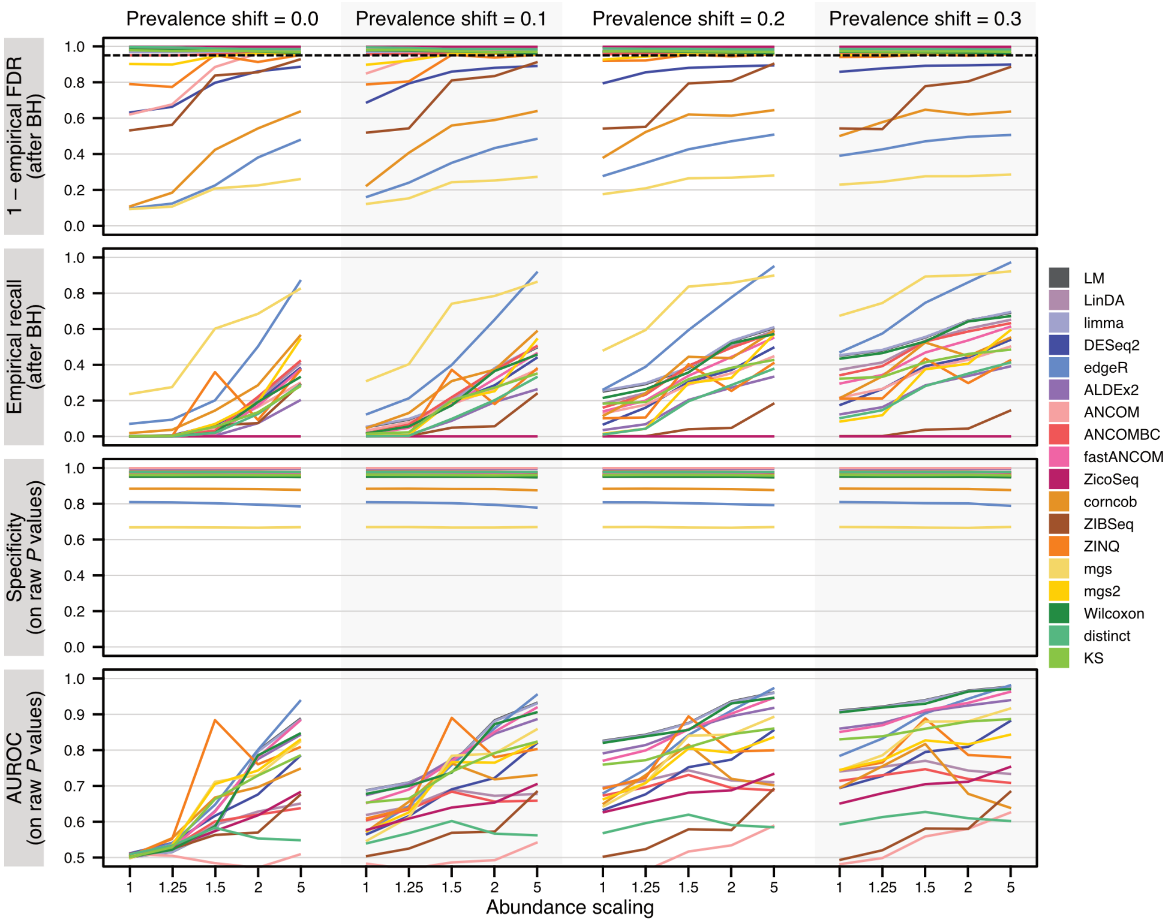


**Supplementary Figure 6: Performance of differential abundance testing methods improves with increasing effect size.** The mean empirical FDR, mean empirical recall, the mean specificity (1 - false positive rate, calculated on raw *P* values) and the mean AUROC (on raw *P* values) for the detection of ground truth DA features are shown across all included methods for varying effect sizes of the same signal implantation benchmark (all features eligible for implantation). All values were recorded for a sample size of 100. With larger effect sizes (both prevalence shifts and scaled abundances), the AUROC for the detection of ground truth DA features generally increased. For ANCOM, lines for mean FDR and recall are dashed, since ANCOM does not output *P* values and multiple testing corrections cannot be applied (see **Methods**).

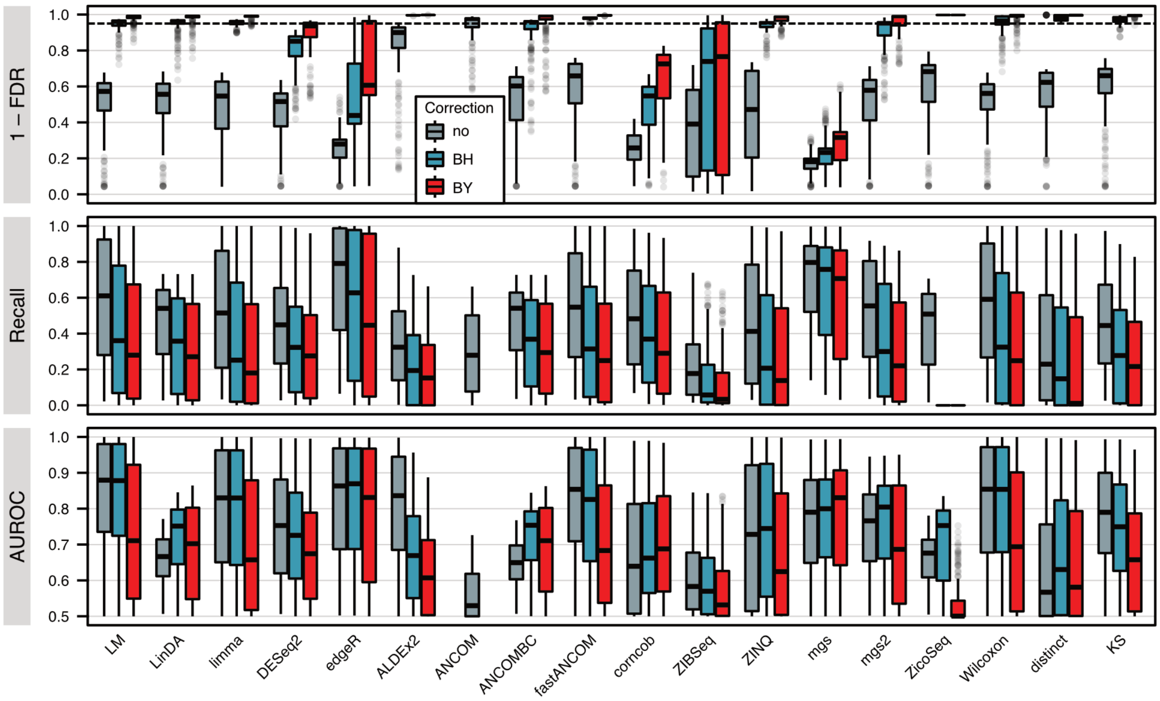


**Supplementary Figure 7: The effect of different adjustment methods for multiple hypothesis testing.** For each included DA testing method, FDR, recall, and AUROC for the detection of differentially abundant features was calculated at a cutoff of 5%, using either the raw *P* values (no correction), or the results of the Benjamini-Hochberg correction (BH), or the Benjamini-Yekutieli correction (BY). Box plots show the distribution of all values across all repetitions and included effect sizes. As expected, a cutoff of 5% on raw *P* values results in high FDR across all methods except for *ALDEx2*, which appears to be overconservative – in line with previous benchmarks[2]. After either BH and BY correction, most methods properly control the empirical FDR at the nominal 5% level, while those methods that failed to do so under BH, also failed under BY (e.g. *metagenomeSeq (mgs)*, *edgeR*, or *corncob*). The BY correction not only results in lower empirical FDR across all methods, but it also generally leads to reduced recall and lower AUROC. See **Fig. 1** in the main text for boxplot definitions. Since ANCOM does not output *P* values (see **Methods**), BH and BY corrections cannot be applied, so the corresponding values cannot be shown.

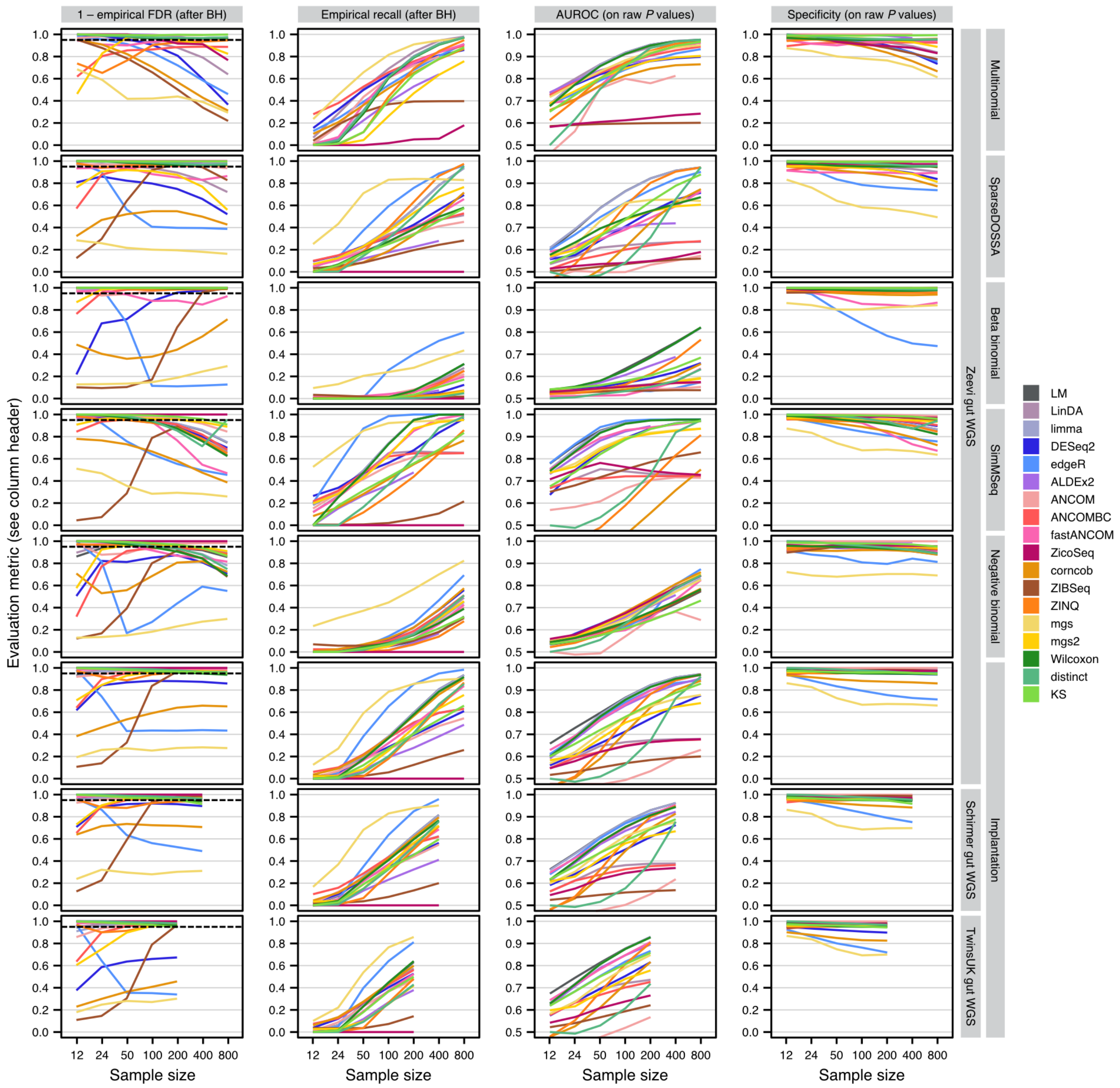


**Supplementary Figure 8: Performance of differential abundance testing methods varies largely across simulation frameworks and comparably little across datasets.** The evaluation results of different underlying simulation frameworks are visually compared across the top 6 rows (see labels to the right) and contrasted to the differences in evaluations resulting from the use of different data sets in the implantation framework across the bottom three rows. For a single, moderate effect size (abundance scaling of 2, prevalence shift of 0.1, if applicable see **Fig. 2**), the mean empirical FDR, mean empirical recall (both computed after BH correction of raw *P* values), the mean specificity, and mean AUROC values for the detection of differentially abundant features are shown across all repetitions for all included DA testing methods (see also **Methods**). Data simulated by parametric simulation frameworks used the real Zeevi gut WGS dataset as input. For ANCOM, lines for empirical FDR and recall are dashed, since ANCOM does not output *P* values (see **Methods**).

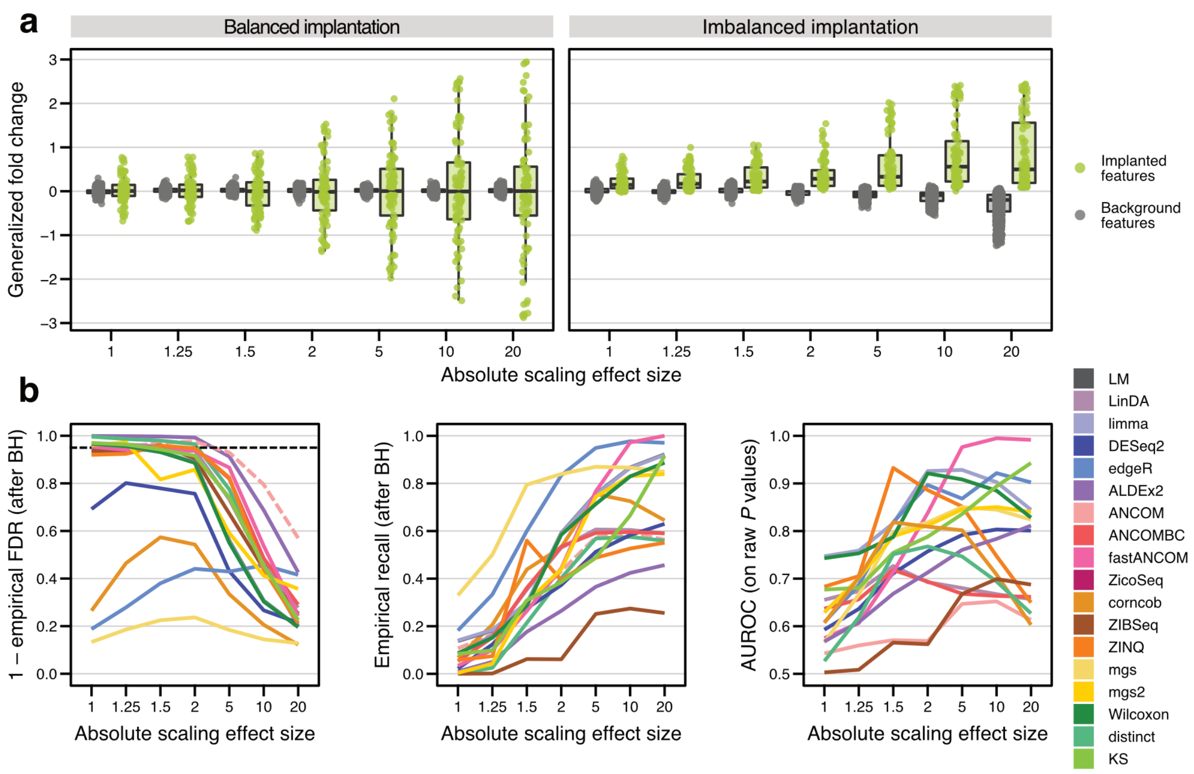


**Supplementary Figure 9: Performance of differential abundance testing methods in the presence of strong compositionality effects. a)** The generalized fold change is shown across different effect sizes for background features and implanted features for the default signal implantation framework with a balanced implantation into both groups (left, see **Methods** and Weiss *et al.*[3]) and the imbalanced implantation (right, see **Methods** and also Jonsson *et al.*[4]). In the default framework, compositional effects are minimized, meaning that background (unimplanted) features show no difference between groups, whereas in the imbalanced implantations, features are only implanted into a single group, thereby leading to abundance shifts in background features (average generalized fold change decreasing with effect size) as a consequence of compositionality. Generalized fold changes for a single representative repetition are shown (prevalence shift of 0.2, only high abundant features eligible for implantation, see **Fig. 1** in the main text for boxplot definition). **b)** The mean empirical FDR, mean empirical recall, and mean AUROC values are shown across all repetitions (prevalence shift of 0.2, only high abundant features eligible for implantation, sample size of 200) for each abundance scaling effect size (with which the compositionality effects also increase) for all included DA methods (see **Methods**). While compositionality (at extreme effect sizes) leads to a general trend of deteriorating DA method performance, *ANCOM* and *ALDEx2* are least affected in terms of FDR. For ANCOM, lines for empirical FDR and recall are dashed, since ANCOM does not output *P* values (see **Methods**).

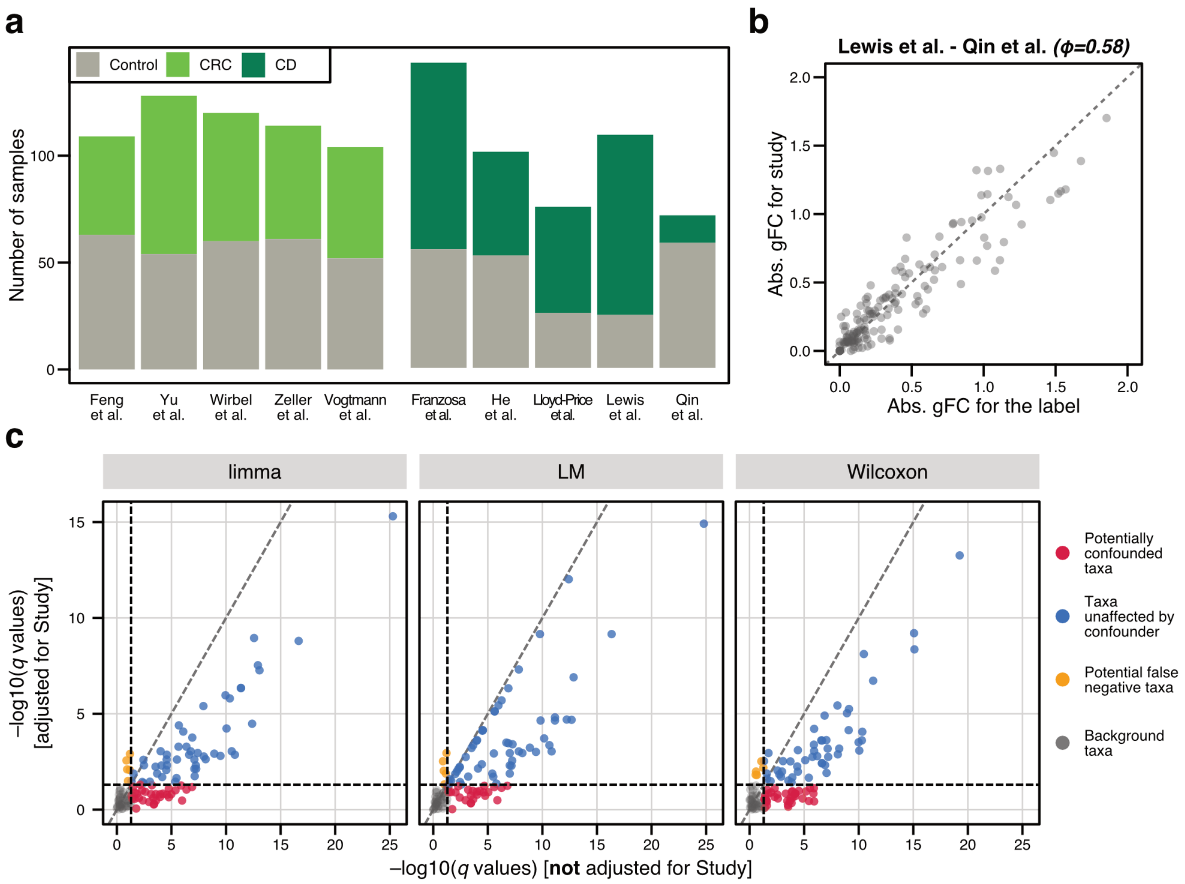


**Supplementary Figure 10: Study heterogeneity in real data resembles broad confounding conditions as simulated using the signal implantation framework. a)** For both colorectal cancer (CRC) and Crohn's disease (CD), the number of samples in each group (control and respective case group) is shown across studies as a bar plot. For CRC, studies are generally balanced, while there are larger differences in group proportions for CD studies. **b)** Across all pairwise combinations of CD studies, Lewis *et al.*[6] and Qin *et al.*[7] exhibit the strongest confounding potential due to study heterogeneity ($\phi$=0.58 between study origin and disease status). For this study combination, the generalized fold change associated with the disease label (CD) and the study origin are plotted against each other across all included bacterial taxa, showing a pronounced correlation. **c)** Using both the naive (not adjusted for study heterogeneity as a confounder) and the study-adjusted configuration of *limma*, the *LM*, and the *Wilcoxon* test, all bacterial genera from the combination of Lewis *et al.* and Qin *et al.* (see b) were tested for differential abundance between control and CD samples. The resulting estimated FDR values (Benjamini-Hochberg corrected *P* values) are plotted against each other as scatter plots, with 0.05 indicated as dashed black line for both models. Red dots highlight taxa that are potentially confounded (identified as differentially abundant in the naive model, but not the confounder-adjusted model), with a similar proportion of potentially confounded taxa across all three DA testing methods. Taxa that are significantly associated with CD, independent of the type of model used, are highlighted in blue, whereas taxa that were not significant in the naive model, but do appear significantly different in the confounder-adjusted model are shown in orange.

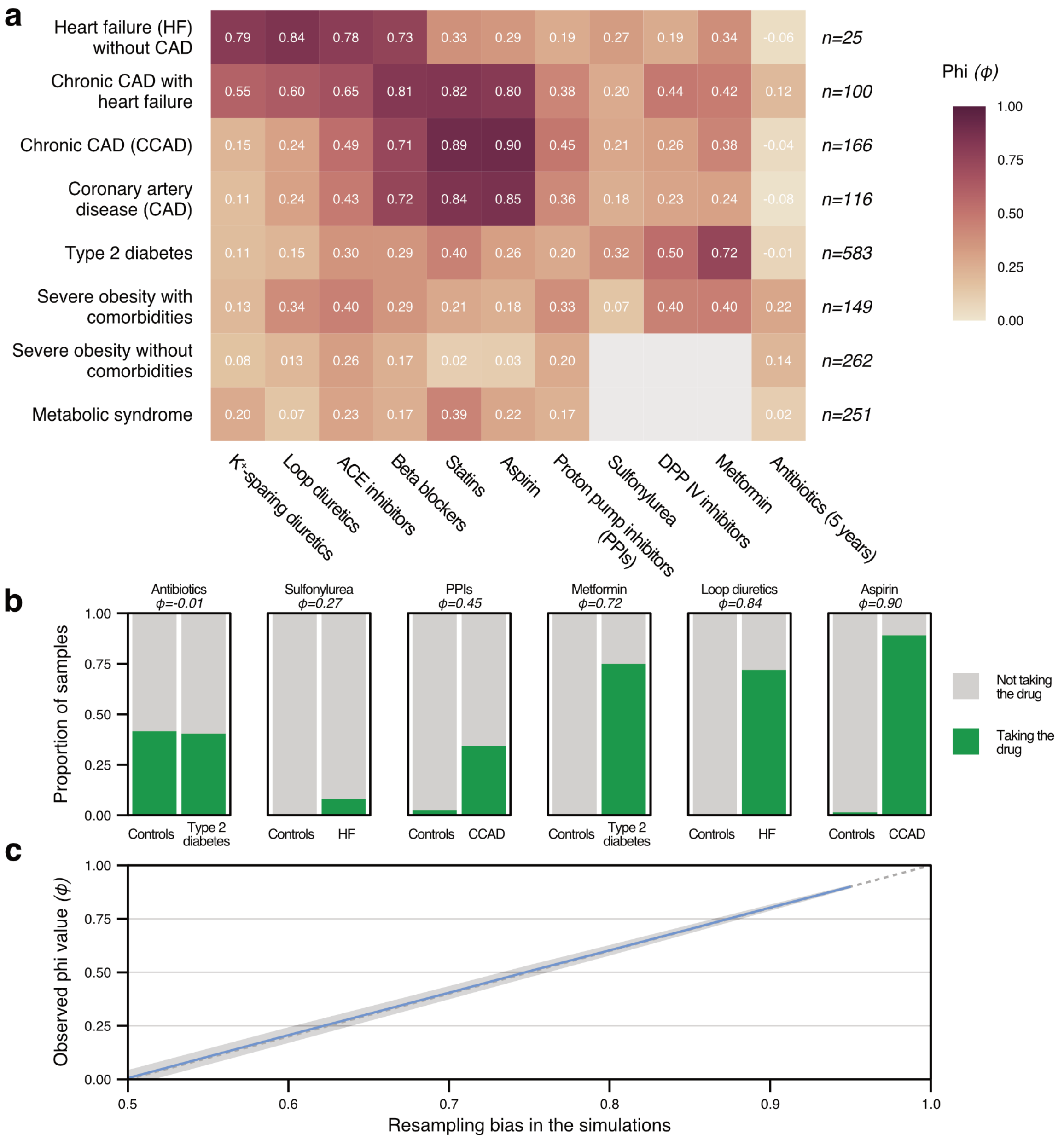


**Supplementary Figure 11: Empirical phi coefficients between cardiometabolic diseases and medication. a)** Phi coefficients were calculated between different disease groups from the MetaCardis cohort[5] and 330 healthy controls (see **Methods**). For a given disease, the highest phi values are observed for the most common drug indications. Gray squares represent NAs, which resulted when no individuals in either case or control group were taking a given medication. **b)** Medication intake broken down by case or control group. High concomitance between drug intake and disease status produces large phi coefficients, while negative coefficients indicate that the control group was more medicated than a given disease cohort. **c)** Linear relationship (with 95% confidence interval as shaded gray area) between the bias parameter used in our framework to produce confounded simulations (see **Methods**) and empirical phi values of simulated data.

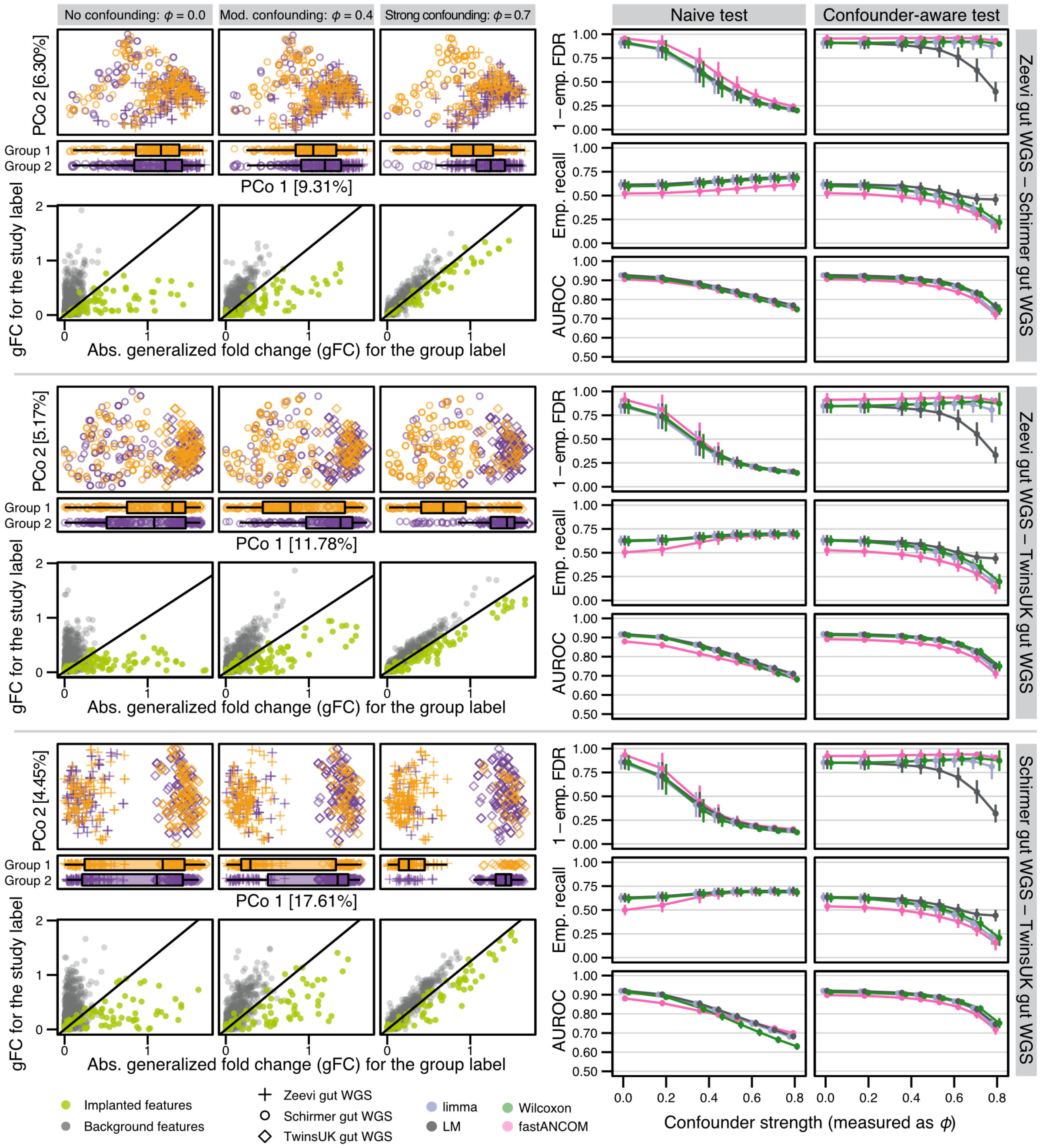


**Supplementary Figure 12: Broad confounding results in slightly worse performance of DA testing methods compared to narrow confounding.** For each two-way combination of the included gut WGS studies (Zeevi WGS, Schirmer WGS, and TwinsUK WGS), implantation simulations were created from data of both studies as input, using the study information as confounding variable for the generation of (biased) resampled testing sets (see **Methods**). On the left, principle coordinate projections for various levels of confounding are shown, visualizing how the study and group variables become more aligned with increased confounding strength (as seen in the group-resolved boxplots for the first principle coordinate). Underneath, the absolute generalized fold change (gFC) for the group label is contrasted with the gFC for the study label, with implanted features highlighted in light green (see **Fig. 3** in the main text for reference). On the right side, the performance of *limma*, the *LM*, *fastANCOM,* and the *Wilcoxon* test are shown in dependence of the confounder strength as measured by ϕ (see **Methods**). Each test was run in the 'naive' (without adjusting for the study variable, left column) and in the confounder-adjusted configuration (right column). Empirical FDR and recall were calculated after Benjamini-Hochberg correction, while AUROC was calculated on raw *P* values.

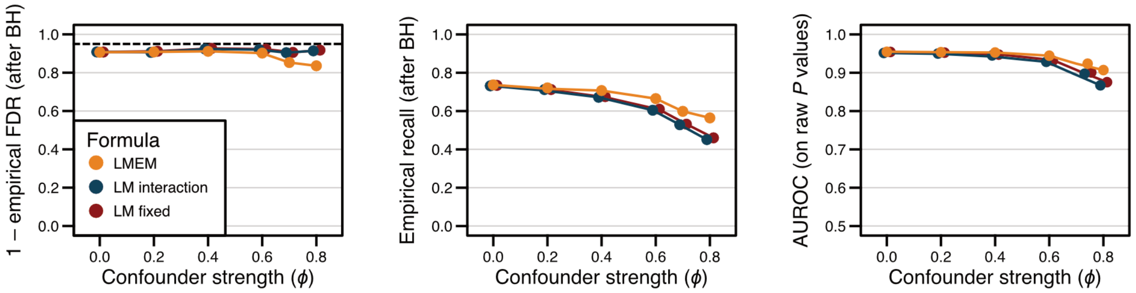


**Supplementary Figure 13: Performance of confounder-adjusted linear models is minimally impacted by choice of model formula.** Different ways of adjusting the linear model for confounders were explored for the same simulation as shown as in **Fig. 3** (abundance scaling factor of 2, prevalence shift of 0.2, all features eligible for implantation, sample size of 200 shown here). LMEM represents the random effect model used in the main text for confounder adjustment, i.e. with formula *lmerTest::lmer(feature~label + (1|confounder)),* which was fit using the base R *summary* function. The LM interaction model had the formula *lm(feature~label * confounder)* and was also fit with the *summary* function. The fixed effect model was implemented as *lm(feature~label + confounder),* and the significance of the label variable was tested using a Type III analysis of variance (via the *car::Anova* function), i.e. one which does not depend on the order of terms in the model formula. The LMEM had a slightly higher proportion of false positives than the other methods at confounder strengths > 0.6, albeit coupled with higher recall and AUROC, but results were largely similar to one another in our evaluation.

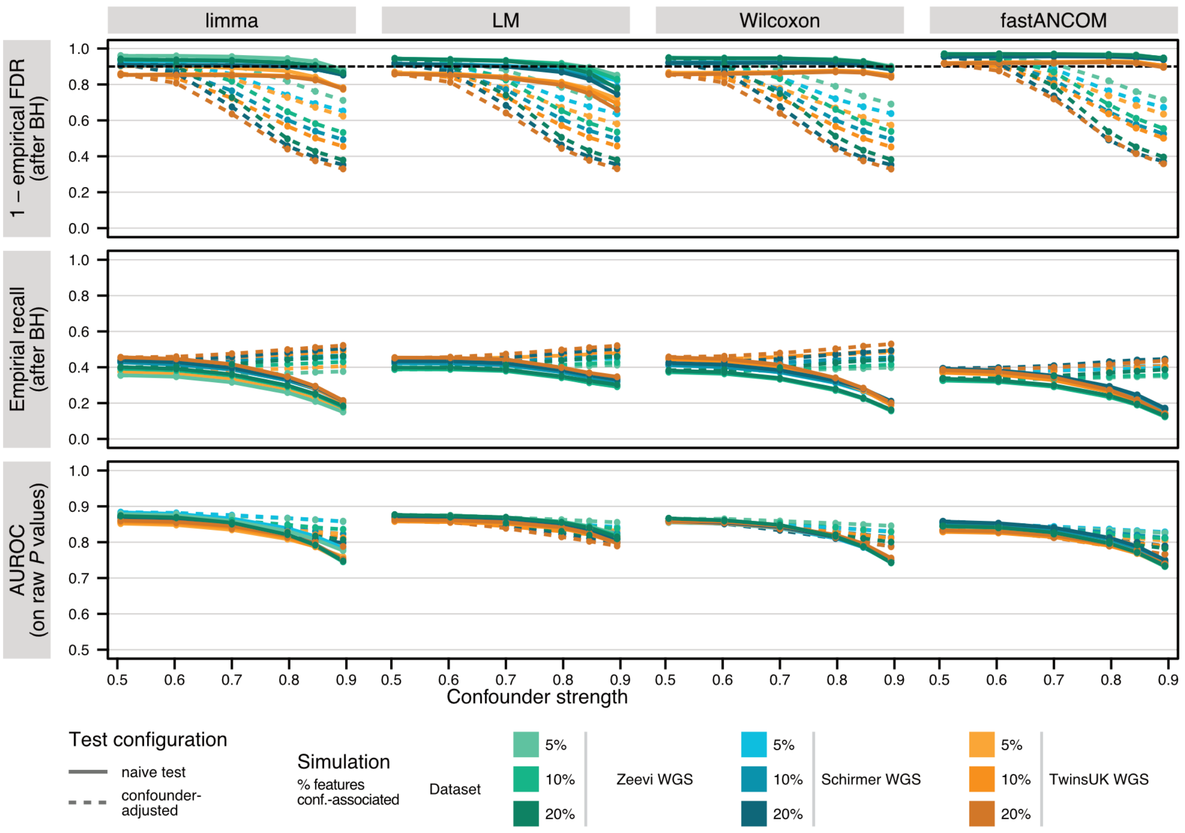


**Supplementary Figure 14: Confounder-adjusted DA testing methods show similar performance across datasets and varying proportion of confounder-associated features.** For all included gut WGS studies (Zeevi WGS, Schirmer WGS, and TwinsUK WGS), several confounded simulations were created with a varying number of features implanted as confounder-associated features into the simulations (5%, 10% or 20% of features implanted with the confounder label, always 10% of features implanted with the main group label, see **Methods**). The mean empirical FDR, the mean empirical recall, and the AUROC values for the detection of differentially abundant features were calculated for each simulation and each of the included DA testing methods (*limma*, the *LM*, the *Wilcoxon* test, and *fastANCOM*), while taking either the confounder variable into account or not (naive and confounder-adjusted models, respectively). Lines show the mean performance of each method across all repetitions for a single effect size (abundance scaling of 2, prevalence shift of 0.2) for each different simulation. All values were recorded for a sample size of 100.

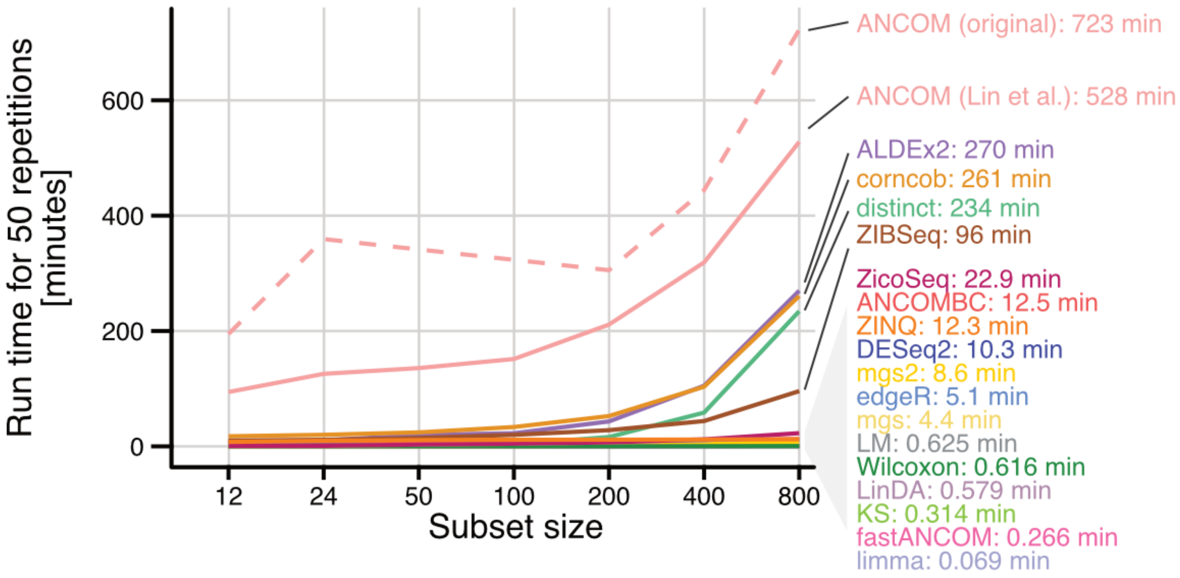


**Supplementary Figure 15: Comparison of runtime across differential abundance testing methods.** Runtime was recorded on the same machine for 50 repetition of different subset sizes from a single repetition in the same signal implantation benchmark (abundance scaling of 2, prevalence shift of 0.1, all features eligible for implantation). Methods are annotated with the time needed to run the subset size of 800 samples on the same laptop. The original *ANCOM* implementation was obtained from the website of the first author of the *ANCOM* manuscript at https://sites.google.com/site/siddharthamandal1985/research.

#### Supplementary Note 1: On the need to make fewer assumptions in large-scale, heterogeneous datasets

Our work was aimed at closing the gap between existing benchmark criteria and the contemporary reality of microbiome data analysis. In designing our simulations, we considered the fundamental task of differential abundance testing of case-control taxonomic profiles. We chose this specific application because it is at the core of most human disease association studies (including those based on 16S rRNA gene markers), and because bacterial taxa are known to vary substantially between individuals (whereas the functional properties exhibited by those taxa collectively are relatively stable, see e.g. Figure 2 from the Human Microbiome Project Consortium[8]). Furthermore, taxonomic profiles in particular have demonstrably different statistical properties than simulated datasets thus far used in DA method benchmarks[3,4,9,10] (see main text **Fig. 1a**), making their recommendations potentially unreliable for real microbiome association studies. A previous benchmark by Hawinkel *et al.*[11] showed that some DA methods report spuriously low *P* values under the null hypothesis being true (especially in sparse taxa with low abundances), suggesting both misapplication and/or method-inherent insufficient type I error control in taxonomic settings. To better characterize this behavior and estimate its impact in applied microbiome studies, we used simulations which we demonstrated to retain key characteristics of several real input datasets (including multiple body sites), and we included additional scenarios (i.e. confounding) and DA methods in our benchmark. Our findings were found to agree with those in Hawinkel *et al.* yet also greatly expanded the scope of inquiry, allowing us to more empirically delineate conditions affecting method performance.

Bulk RNA-seq methods were originally developed for few biological replicates (in contrast to the scope and scale of current microbiome research[12,13]), and test the null hypothesis that features have the same mean under two conditions (which is not robust to outliers or high between-sample variance characteristic of taxonomic microbiome profiles). To test their suitability in a large-N setting, Li *et al.*[14] use several real datasets from the Genotype-Tissue Expression (GTEx[15]) and the Cancer Genome Atlas (TCGA[16]) and show that tools like *DESeq2* and *edgeR* display exaggerated false positives in population-level RNA-seq studies (containing fifty to hundreds of samples per group). These authors also find the *Wilcoxon* test's more conservative, rank-based hypothesis to empirically outperform other methods, especially in terms of FDR control (verified by permutation). They conclude that parametric methods should only be used when the per-condition sample size is less than eight and power is a concern.

Our related conclusion that restrictive parametric methods do not offer advantages over classical statistical methods when applied to human-associated bacterial profiles – and even have negative consequences – contradicts the results from previous microbiome-specific benchmarks. These discrepancies can be explained by differences in the design of the underlying simulation strategies. Jonsson *et al.*[4] concluded *DESeq2* and *edgeR* to have the best overall performance – probably owing to their small sample sizes (two groups of 3, 6 and 10 vs. two groups of 6, 12, 25, 50 100, 200, 400, and 800 here) and exclusive use of gene-level abundances, which are well-established[8] to display far lower between-sample variance than the sparse taxonomic profiles investigated here. Reaching a similar conclusion, McMurdie and Holmes[9] explored two groups of 3, 5, and 10, and used parametric multinomial simulations, which insufficiently capture relevant statistical properties of taxonomic profiles (see main text **Fig. 1a-c**). As we demonstrate, their conclusions are not supported when larger sample sizes and more realistic simulation procedures are considered (see **SFig. 8**).

#### Supplementary Note 2: Theoretical considerations for calibrating different methods

In our benchmark we observed a high empirical false positive rate for several methods independent of the multiple hypothesis correction procedure used (Benjamini-Hochberg or Benjamini-Yekutieli, see **SFig. 7**), suggesting method-inherent issues with type I error control. One obvious culprit for this is of course unmet distributional assumptions embedded in the DA models, but another possible explanation for these high empirical FDR values could be that some methods are not well-calibrated for microbiome data and report universally low *P* values, while still being able to give lower *P* values to implanted features compared to background features – in essence, correctly ranking ground truth bacterial taxa. Such cases would result in high AUROC values on uncorrected *P* values, and performance could theoretically be improved by changing the *P* value cutoff for significance. However, of those methods with high empirical FDR, only *edgeR* showed comparably high AUROC values (see main text **Fig. 2**) and therefore fit this pattern, whereas other methods (such as *metagenomeSeq* or *corncob*) could not even theoretically be recalibrated to high precision and high recall.
